# Human pluripotent stem cell-derived respiratory airway progenitors generate alveolar epithelial cells and recapitulate features of idiopathic pulmonary fibrosis

**DOI:** 10.1101/2023.01.30.526265

**Authors:** Mikael G. Pezet, Juan A. Torres, Tania A. Thimraj, Ivana Matkovic Leko, Nadine Schrode, John W. Murray, Kristin G. Beaumont, Hans-Willem Snoeck

## Abstract

Human lungs contain unique cell populations in distal respiratory airways (RAs). These populations accumulate in patients with lung injury, chronic obstructive pulmonary disease (COPD) and idiopathic pulmonary fibrosis (IPF). Their lineage potentials and roles are unknown, however. As they are absent in rodents, deeper understanding of these cells requires a human *in vitro* model. Here we report the generation from human pluripotent stem cells (hPSCs) of expandable spheres (‘induced respiratory airway progenitors’ (iRAPs)) consisting of all RA-associated cell types. iRAPs could differentiate into type 1 (AT1) and type 2 alveolar (AT2) epithelial cells in defined conditions, showing that alveolar cells can be derived from RAs. iRAPs with deletion of HPS1, which causes pulmonary fibrosis in humans, display defects that are hallmarks of IPF, indicating involvement of intrinsic dysfunction of RA-associated cells in IPF. iRAPs thus provide a model to gain insight into human lung regeneration and into pathogenesis of IPF.

## INTRODUCTION

The respiratory epithelium consists of basal (BC), ciliated, secretory, goblet and neuroendocrine cells in the airways, and alveolar type 1 (AT1) and surfactant-producing alveolar type 2 (AT2) cells in the alveoli.^1^ AT1 cells, through which gas exchange takes place, are the primary cell of the respiratory system, and lung regeneration after injury is geared towards restoring AT1 function. The best characterized mechanism is the conversion of a fraction of AT2 cells to AT1 cells through transitional intermediates,^2–4^ although AT1-to-AT2 plasticity exists in the neonatal mouse lung.^5^ While most mouse studies on alveolar repair after injury are based on the premise of AT2-to-AT1 conversion, lineage tracing studies have suggested a contribution of cells with characteristics of airway secretory cells after severe injury.^2,6–11^ Multiple cellular pathways might therefore converge on alveolar repair.

Putative regenerative cells with features of airway secretory cells have recently also been identified in humans. In contrast to mice, humans, primates and ferrets have ‘respiratory airways’ (RAs) or ‘terminal respiratory bronchioles’ (TRBs), representing the most distal branches of airway tree into which individual alveoli open and that terminate in alveolar ducts, where multiple alveoli coalesce. Expression of the secretory cell marker, *SCGB3A2*, was identified in scRNA-seq studies of the distal lung in cells also co-expressing AT1 and AT2 markers.^12^ Basil et al. reported presence of *SCGB3A2^+^*cells negative for the AT2 marker, *SFPTC*, in RAs.^13^ Murthy et al. identified multiple subpopulations in RAs characterized by co-expression of another AT2 marker, *SFTPB* (**Fig. 1a**).^14^ The increased abundance of these populations after lung injury, in COPD and in IPF suggests a role in regeneration.^12–14^ We therefore call these cells ‘respiratory airway progenitors’ (RAPs). In addition to RAPs, IPF lungs also show accumulation of ‘aberrant basaloid cells’ (aBCs), a population that is much less abundant, though not absent, in normal lungs and expresses *KRT17*, *COL1A1* and other extracellular matrix (ECM) components but lacks common BC markers such as *KRT5*.^12,15^

**Figure 1.**
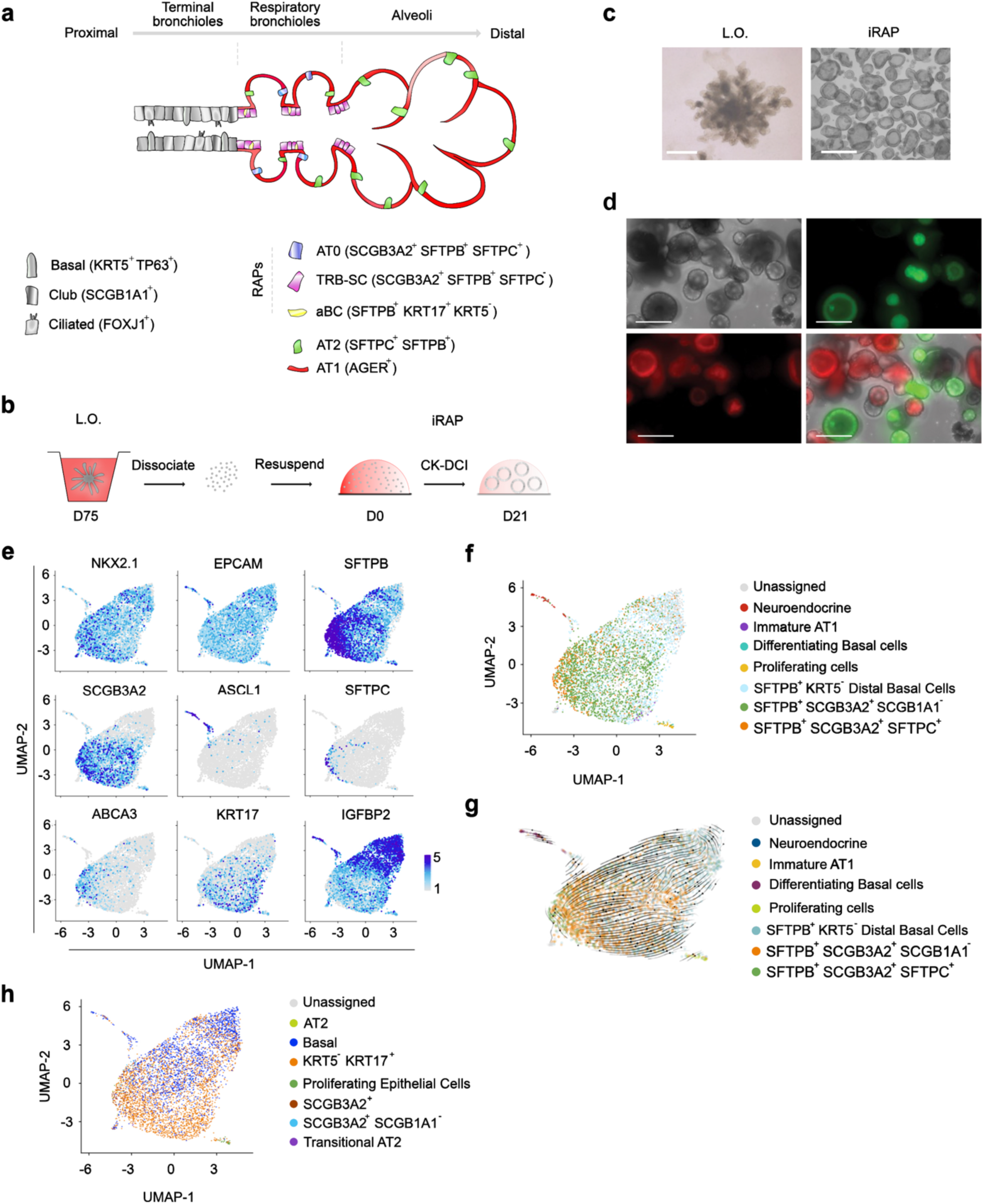
ScRNA-seq analysis of iRAPs. **a.** Schematic of the anatomical arrangement and markers of RAPs *in vivo*, based on ref. 14. **b.** Schematic of experimental design to generate iRAPs from LOs**. c.** Representative bright field images of iRAPs generated from ESCs. **d.** iRAPs generated from mixed LO single cells expressing either GFP or mScarlet. **e.** Feature plots for indicated markers. **f.** Cell identity assignment based on data from ref. 14. **g.** scVelo latent time velocity analysis. **h.** Cell identity assignment based on data of ref. 12 with inclusion of data from IPF lungs.

As they are absent in rodents, insight into the lineage potentials and role of RAPs requires a human *in vitro* model. Here we report the generation from human pluripotent stem cells (hPSCs) of expandable, clonal spheres that consist of all cell populations present in RAs and can differentiate into AT2 and AT1 cells. These ‘induced respiratory airway progenitors’ (iRAPs) display hallmarks of IPF when carrying a mutation associated with familial IPF, thus suggesting a link between intrinsic defects of RAPs and IPF pathogenesis. iRAPs thus provide an *in vitro* model to gain insight into mechanisms underlying normal and pathological human alveolar regeneration and to discover and validate drug targets.

### Generation of transitional lung spheres

We previously developed a directed differentiation strategy of hPSCs (embryonic (ESCs) and induced pluripotent stem cells (iPSC)) into 3D lung organoids (LOs) that undergo a process similar to branching morphogenesis.^16,17^ We cultured dissociated cells from LOs in medium supplemented with CHIR (GSK3 inhibitor), keratinocyte growth factor (KGF), dexamethasone, 3′,5′-cyclic adenosine monophosphate (cAMP), and 3-isobutyl-1-methylxanthine (IBMX) (CK-DCI) (**Fig. 1b**), previously reported to promote outgrowth of spheres containing AT2 cells (alveolospheres) from hPSC-derived lung progenitors.^18–20^ Within 2 weeks, hollow spheres developed (**ESC Fig.1c; iPSC ED1a**) that could be serially passaged every ∼3 weeks for up to at least 7 passages with an average expansion of ∼10-fold between each passage **(ESC ED2a, iPSC ED1b)**. Generation of spheres from mixtures of LO cells expressing either GFP or mScarlet yielded spheres with only one fluorescent reporter, indicative of clonality (**Fig. 1d**).

scRNA-seq revealed a relatively homogenous population with respect to expression of epithelial (*EPCAM)*, lung (*NKX2.1)*, and AT2/RAP (*SFTPB)* markers (**Fig. 1e**). A large subpopulation expressed *SCGB3A2*, with some *SCGB3A2^+^* cells co-expressing AT2 markers (*SFTPC*, *NAPSA* and *ABCA3)* (**Fig. 1e**). The *SCGB3A2^-^* population expressed *IGFBP2*, a BC marker according to LungMap (**Fig. 1e**).^21^ Murthy et al. described several subpopulations residing in human terminal airways, including *SFPTB*^+^*SCGB3A2^-^*distal BCs, some of which are low to negative for the airway BC markers, *KRT5* and *p63*, *SFTPB*^+^S*CGB3A2*^+^*SCGB1A1*^+^ pre-TRB secretory cells (pre-TRB-SCs), *SFTPB*^+^S*CGB3A2*^+^*SFTPC*^-^ TRB-SCs, and, finally, *SCGB3A2*^+^*SFTPB*^+^*SFTPC*^+^ “AT0” cells near the alveolar sacs (**Fig. 1a**).^14^ Cell type annotation using machine learning trained on these data showed that the *SCGB3A2^+^* population consisted of AT0 cells and TRB-SCs, whereas the *SCGB3A2^-^IGPBP2^+^*population mapped to distal BCs (**Fig. 1f**). Interspersed between those were ‘differentiating BCs’ and rare ‘immature AT1 cells’, likely representing differentiation intermediates. Except for a small, distinct population of neuroendocrine cells, no match with any other lung populations was found. Because the spheres contained almost exclusively cells associated with RAs, we called these cells ‘induced respiratory airways progenitors’ (iRAPs).

Trajectory analysis using scVelo^22^ showed differentiation pathways originating from distal BCs and terminating in AT0 cells (**Fig. 1g**). These findings suggest a lineage from distal BCs over TRB-SCs to AT0 cells that was not recognized in recent scRNA-seq studies of human lung but matches the proximodistal arrangement of these cell types *in vivo* (**Fig. 1a**). Cell identity assignment based on the data of Habermann et al.,^12^ which included samples from IPF patients, showed comparable cell identity assignment while also revealing the presence of *KRT5*^-^*KRT17*^+^ aBC-like cells (**Fig. 1h**)

RT-qPCR for *SFTPC*, *SFTPB* and *SCGB3A2* mRNA confirmed the scRNA-seq analysis **(ESC,** **Fig. 2a; iPSC, ED1c)**. Confocal microscopy revealed uniform expression of EPCAM, NKX2.1 and SFTPB, with pro-SFTPC expressed in some of the spheres **(ESC,** **Fig. 2b; iPSC, ED1d-e)**. SFTPB and SCGB3A2 were expressed within the same spheres **(ED2b)**. Western Blot showed detectable presence of fully processed SFTPB and SFTPC **(ESC** **Fig 2c, iPSC ED1f)**. HT2-280, a marker of mature AT2 cells,^23^ accounted for ∼2% of the cells **(ESC** **Fig. 2d; iPSC ED1g,** representative gating strategy **ED3**). iRAP spheres showed apical accumulation of the lysosomal dye, LysoTracker Red (**ED2c**) and lamellar bodies (LBs) by transmission electron microscopy (**Fig. 2e, arrows**), a feature of AT2 cells. However, these had low electron density and were often organized as multivesicular bodies, indicative of immaturity.^24^ iRAPs maintained a stable phenotype over several passages after passage one (**ED2d**) and were phenotypically similar whether generated from early or late stage LOs (**ED2e**). They can be cryopreserved and thawed while retaining their expression profile (**ED2f**). Together, these observations indicate that iRAP spheres contain a mixture of AT0 cells, TRB-SCs, distal BCs and aBCs, thus encompassing recently identified RAPs in the human distal lung.

**Figure 2.**
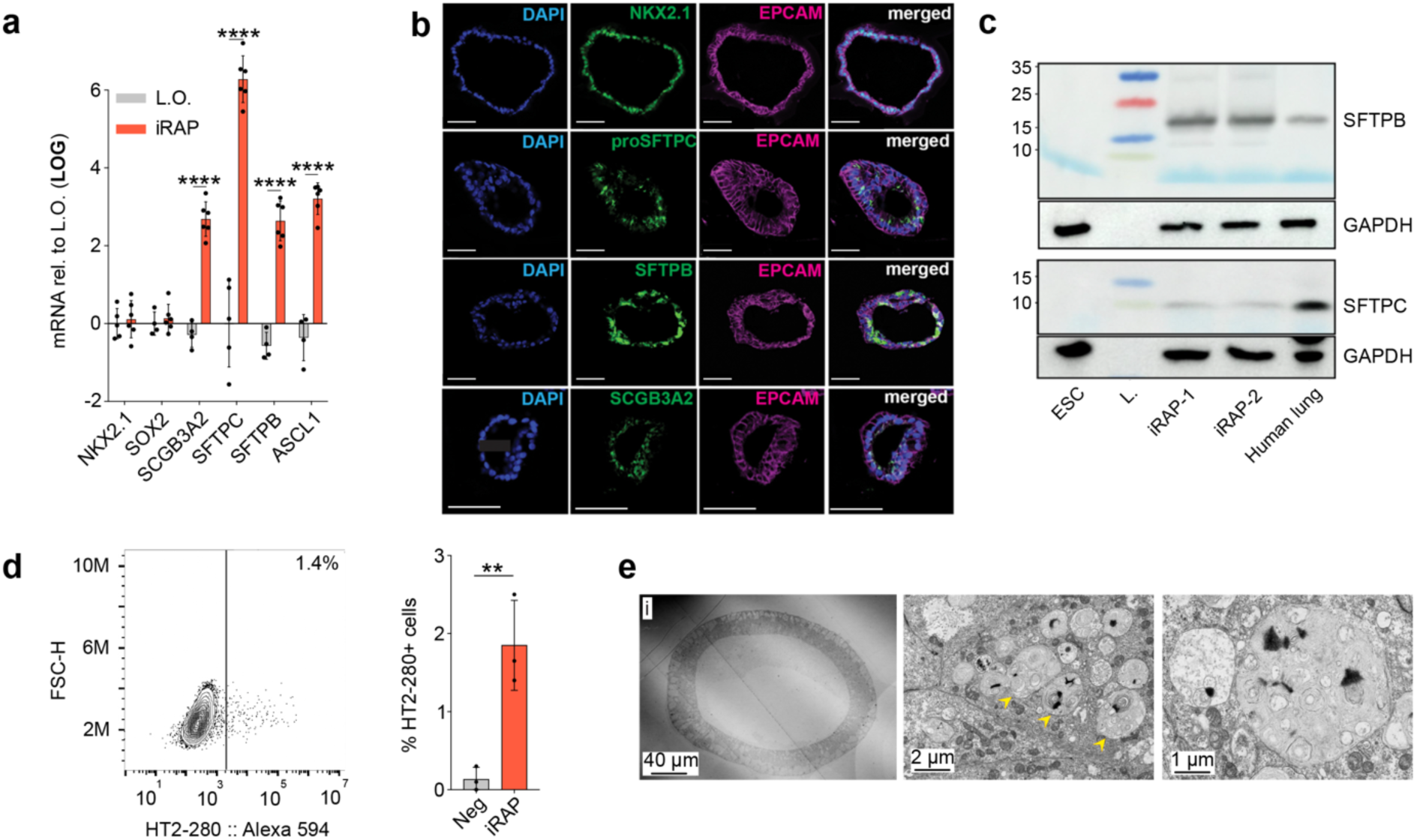
Characterization if iRAPs. **a.** mRNA expression of indicated markers in iRAPs compared to LOs. ****p<0.0001, two-way ANOVA with Sidak multiple comparison test, n=6. **b.** Representative confocal images after staining for indicated markers in iRAPs. Scale bar = 50 µm. **c.** Representative WB for processed SFTPB and SFTPC in ESCs, in two ESC-derived iRAP lines, and in human adult lung. **d.** Representative flow cytometry plot for HT2-280 in iRAPs (left) with quantification (right). **p<0.01, student’s unpaired t-test, n=3. **e.** Transmission electron microscopy of ESC-derived iRAPs (i = 40x, ii = 2000x, and iii = 4000x, arrows: LBs).

### Distal specification of iRAPs

We sought to mature iRAP cells into more distal fates by retraction of CHIR, as previous reports showed that CHIR removal induced AT2 maturation of hPSC-derived alveolospheres.^18,19^ CHIR removal led to disintegration of the spheres (**ED4a),** decreased proliferation (**ED4b)** and a modest increase of the HT2-280^+^ fraction (**ED4c**) which was, however, not corroborated by mRNA expression of the AT2 marker, *SFTPC* (**ED4d**). Furthermore, blocking NOTCH signaling using DAPT, shown previously to induce SFTPC expression in SCGB3A2^+^ cells,^13^ did not significantly affect expression of AT2 markers, except for a modest increase in *ABCA3* mRNA (**ED4e-f).**

To gain deeper insight into signaling pathways driving distal differentiation, we took advantage of publicly available scRNA-seq data (https://fetal-lung.cellgeni.sanger.ac.uk/) collected from human embryonic lung at ∼16-22 weeks post-conception.^25^ Our re-analysis confirmed the presence of a ‘lung epithelium’ cluster (**ED5a**), containing cells expressing *EPCAM*, *NKX2.1*, *SFTPB*, *SFTPC*, *AGER* and *SCGB3A2* (**ED5b**). This cluster contained 5 distinct subclusters (**ED5c**), two of which were of particular interest: the ‘SFTPB^+^SCGB3A2^+^’ cluster, and the ‘AT1-AT2 progenitor’ cluster expressing *AGER*, *SFTPB* and *SFTPC* but no *SCGB3A*2 (**ED5d**). Differential gene expression (DEG) analysis showed that genes defining the ‘AT1-AT2 progenitor’ cluster were affiliated with ‘lung’ and ‘fetal lung’ (**ED5e**). Gene Ontology (GO) analysis suggested involvement of TGF-β signaling (**ED5f**) at this stage of development. UMAP feature plots showed increased expression of SMAD6 and SMAD7, negative regulators of TGF-β signaling, and of NOTUM, a negative regulator of WNT signaling previously shown to be involved in alveolar development (**ED5g)**.^26^ These data suggested that TGF-β and WNT signaling are downregulated during distal differentiation of *SCGB3A2^+^* cells.

To examine whether TGF-β inhibition in the absence of WNT signaling would induce distal differentiation, we cultured iRAPs for 14 days in the presence of DCI and SB-431542, a TGF-β inhibitor^27,28^ (DCI-SB, **Fig. 3a,b**). While the AT1 marker, HT1-56,^29^ is absent in iRAPs cultured in CK-DCI and expression of HT2-280 is low, iRAPs cultured with DCI-SB contained ∼40% HT1-56^+^ and ∼30% HT2-280^+^ cells **(ESC** **Fig. 3c** **and ED6a, iPSC ED1h-j)**. We therefore call the cells distalized iRAPs (D-iRAPs). As no induction of distal markers was observed with DCI or SB alone **(Fig ED6b-e)**, both are necessary for induction of distal fates. BMP inhibition with NOGGIN did not result in induction of HT1-56 or HT2-280, indicating that iRAP differentiation is specifically driven by inhibition of TGF-β (**Fig. 3c**). FGF7 and FG10 are dispensable, and even reduced HT2-280 expression (**Fig. 3c**). RT-qPCR showed increased expression of *SCGB3A2, SFTPC*, *SFTPB* and *AGER* mRNA in D-iRAPs compared to iRAPs (**Fig. 3d**), indicative of a more distal fate. Co-staining for pro-SFTPC and RAGE (the protein encoded by *AGER*) showed that ∼27% of spheres were proSFTPC^+^, 10% were AGER^+^ and 68% were proSFTPC^+^RAGE^+^ (**Fig. 3e,f, ED6f,i**). RAGE and SFTPC were present within the same sphere in 43%, whereas spheres only staining for SFTPC or RAGE represented 27% and 21% of the total, respectively **(Fig. 3e,f, ED6g,h,i)**. scRNA-seq analysis (**Fig. 3g,h**) showed that a large fraction of D-iRAPs expressed *SFTPC*, whereas the highest *SCGB3A2* expression was now found in a population expressing the AT1 markers, *AGER, CLIC5* and *CAV1*, markers that were absent from iRAPs. *ASCL1^+^NEUROD^+^*neuroendocrine cells were still present, while *IGFBP2^+^* distal BCs became rare (**Fig. 3h**).

**Figure 3.**
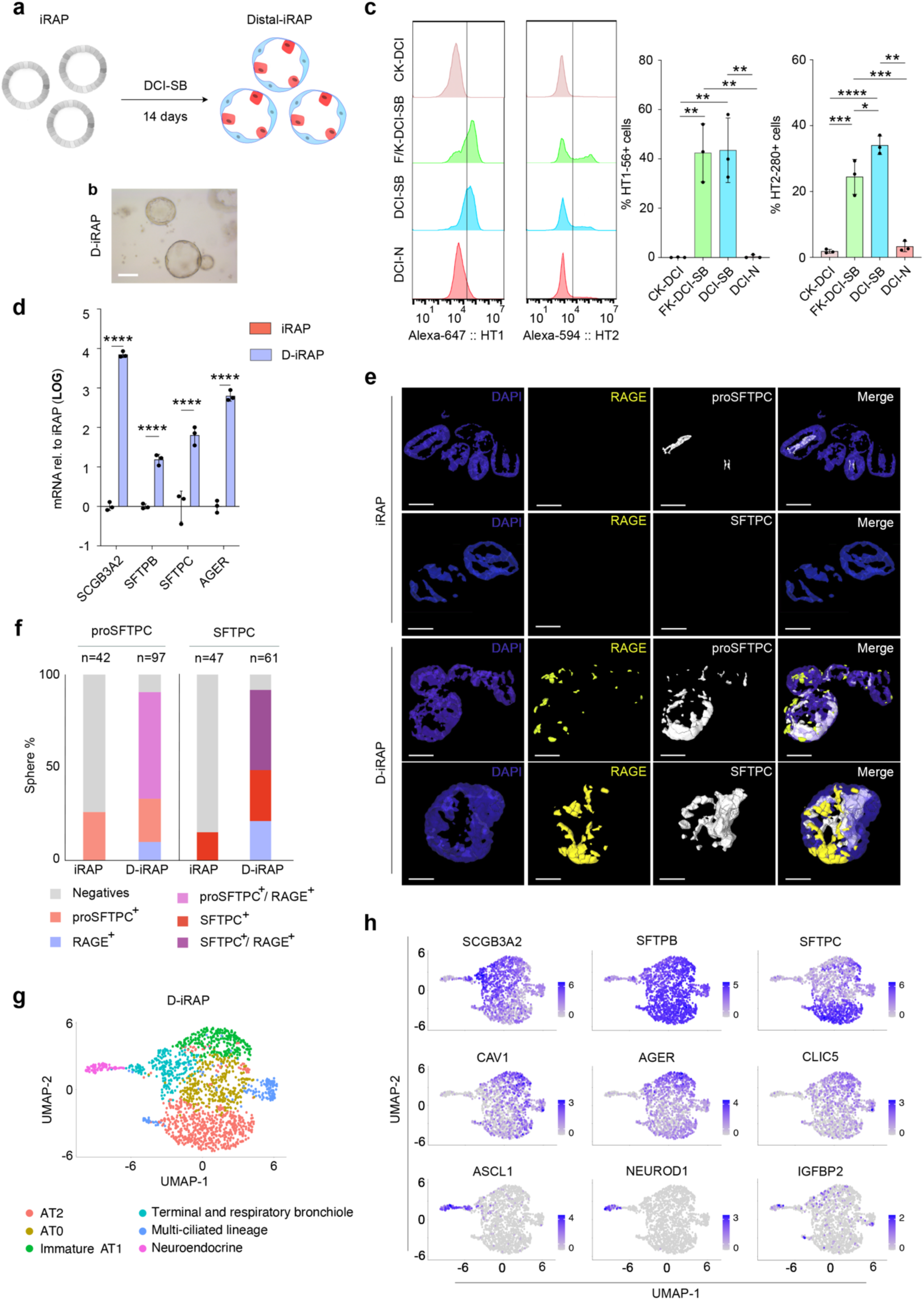
Generation of D-iRAPs. **a.** Schematic of experimental design. **b.** Representative bright field image of D-iRAPs. Scale bar = 200 µm. **c.** Representative flow cytometry histograms (left) and the quantification (right) of expression of HT1-56 and HT2-280 in indicated conditions. *p<0.05, **p<0.01, ***p<0.001, ****p<0.0001, one-way ANOVA with Tukey multiple comparison test, n=3. **d.** Expression of mRNA for the indicated markers in iRAPs and D-iRAPs. **** p<0.0001 two-way ANOVA with Sidak multiple comparison test, n=3. **e.** Z-stack projection of representative 3D confocal images after staining indicated markers. Scale bars = 100 µm. **f.** Quantification of (**e**) in D-iRAPs (pooled from 4 independent replicates). **g.** Cell identity assignment in D-iRAP scRNA-seq dataset based on markers shown in **(h)**. **h.** UMAP feature plots of indicated markers.

We conclude that removal of WNT signaling and inhibition of TGF-β signaling specifies iRAPs toward alveolar epithelial cells.

### Specification of AT1 cells

Expression of AT1 markers and *SCGB3A2* overlapped in D-iRAPs, suggesting incomplete maturation (**Fig. 3h**). A longer differentiation time in DCI-SB conditions up to 21 days did not enhance maturation but decreased cell viability (**ED6j**). We therefore explored other strategies to obtain mature AT1 cells. Physical forces affect AT1 development, suggesting that their specification may require adherent 2D culture conditions.^5,30,31^ Moreover, BMP signaling was previously shown to increase AT1 differentiation efficiency following pneumonectomy in mice,^32^ a finding consistent with our observation that the BMP inhibitor, NOGGIN, negatively impacted the specification of alveolar cell types in iRAPs (**Fig. 3c**). We therefore dissociated D-iRAP spheres and plated the cells on Matrigel-coated plates and subsequently in air-liquid interphase (ALI) cultures. (**Fig. 4a, ED7a**). All cultures were supplemented with ROCK inhibitor which was shown to promote proliferation and survival of airway cells in 2D cultures.^33^ While medium supplemented with BMP4 did not allow attachment of the cells (**ED7b**), high expression of RAGE protein and *AGER* mRNA was maintained in the presence of DCI-SB, SB only and BMP4-SB in submerged 2D culture (**ED7c-d**). Interestingly, the combination of BMP4 and SB downregulated SCGB3A2 and SFTPC protein and mRNA (**ED7c-d**), indicative of AT1 at the expense of AT2 maturation. Similarly, in air liquid interface (ALI) conditions, cells cultured in BMP4-SB maintained confluency (**ED7e-f)**, *AGER*/RAGE (**Fig. 4b-d**) and HT1-56 (**Fig. 4e**) expression but lost expression of *SFTPC* and *SCGB3A2* mRNA (**Fig. 4b**) and protein (**Fig. 4c-d**). More than 95% of the cells expressed HT1-56, whereas HT2-280 became undetectable (**Fig. 4e**). Some *SFTPB* mRNA and protein were still detected however **(ED7g)**. Co-localization analysis showed that SFTPB^+^ cells only accounted for ∼6% of RAGE staining, indicative of an overall mature AT1 lineage (**ED7h,i).** ScRNA-seq analysis showed that >90% of the cells were classified as AT1 cells (**Fig. 4f**). We call these cells induced AT1 (iAT1) cells. A minor fraction was identified as AT2, whereas a small cluster co-expressed *SCGB3A2* and *SFTPC*, indicative of AT0 identity (although query of the Human Lung Cell Atlas^34^ spuriously classified the cells as ‘secretory’ and ‘macrophages’) (**Fig. 4f**). Taken together, iAT1 maturation requires inhibition of TGF-β and activation of BMP signaling in both submerged and ALI 2D conditions.

**Figure 4.**
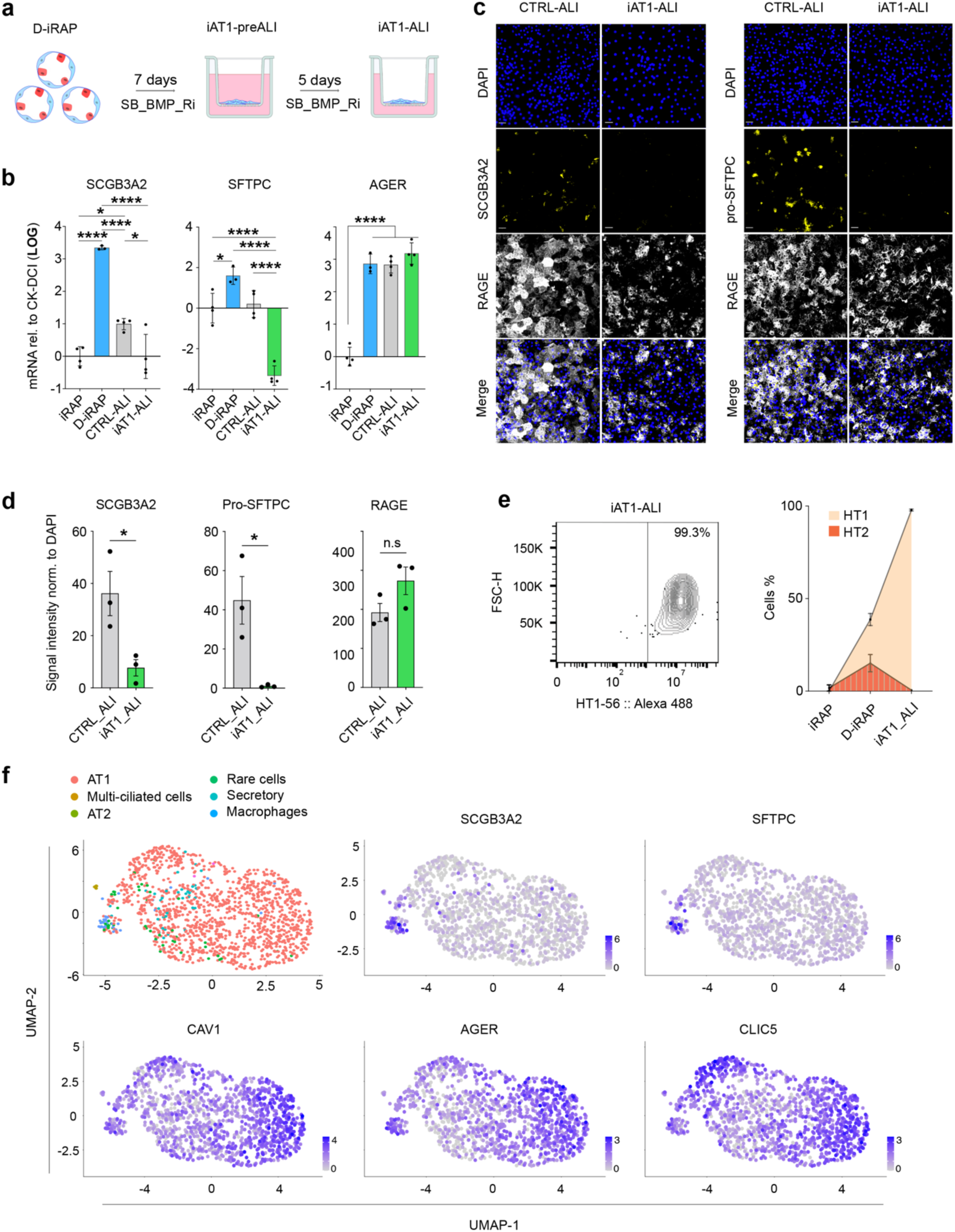
Generation of iAT1 cells. **a.** Schematic of experimental design. **b.** mRNA expression for the indicated markers in various conditions. *p<0.05, ****p<0.0001 One-way ANOVA with Tukey multiple comparison test, n≥4. **c.** Representative confocal images after staining for indicated markers in ALI supported by SB-BMP (iAT1) or DCI-SB (CTRL). Scale bars = 50 µm. **d.** Quantification of (**c**). *p<0.05, Student’s unpaired t-test, n≥3. **e**. Representative flow cytometry plot for HT1-56 (left) and quantification for HT1-56 and HT2-280 (right), n=3. **f.** Cell identity assignment based on Human Lung Cell Atlas of iAT1 scRNA-seq dataset (top left) with UMAP feature plots of indicated markers.

Merged analysis of scRNA-seq data from the iRAPs, D-iRAPs and iAT1 cells (**Fig. 5a**) confirmed acquisition of the AT1-AT2 markers during the transition from iRAPs to D-iRAPs and maturation of AT1 cells with loss of *SFTPC* and *SCGB3A2* in the iAT1 stage, where *AGER^+^CAV1^+^*cells are predominant. (**Fig. 5b-d**). Proximal lung and digestive tract markers were absent at each stage of the development (**Fig. 5d**). Bulk RNAseq comparing cells at the iRAP, D-iRAP and iAT1 stage of the differentiation protocol (**Fig. 5a**) led to similar conclusions **(Fig. 5e,f; ED8a)**. There were 2909 differentially expressed genes between iRAPs and iAT1 cells, and 2821 between D-iRAPs and iAT1 cells stages (**ED8b**, **Source Data 2**). Gene ontology (GO) analysis showed significant induction of the terms, ‘cell shape’, ‘actin filaments’ and ‘cell junctions’ consistent with extensive remodeling of cell morphology during AT1 differentiation **(Fig. 5g; ED8c-d).** Furthermore, the ‘HIPPO signaling’ pathway, which is essential for the generation of AT1 cells during development and after injury in mice,^35–37^ was a significantly induced pathway (**Fig. 5g,h**). Interaction analysis using String-DB^38^ showed that DEGs involved in regulation of actomyosin cytoskeleton and cell junctions were connected to Hippo signaling **(Fig. 5i, ED8e)**, consistent with the reported crosstalk between small GTPase and Hippo signaling^39^ and with the role of CDC42 in epithelial morphogenesis^40^ and acquisition and maintenance of AT1 identity.^5,31,35^

**Figure 5.**
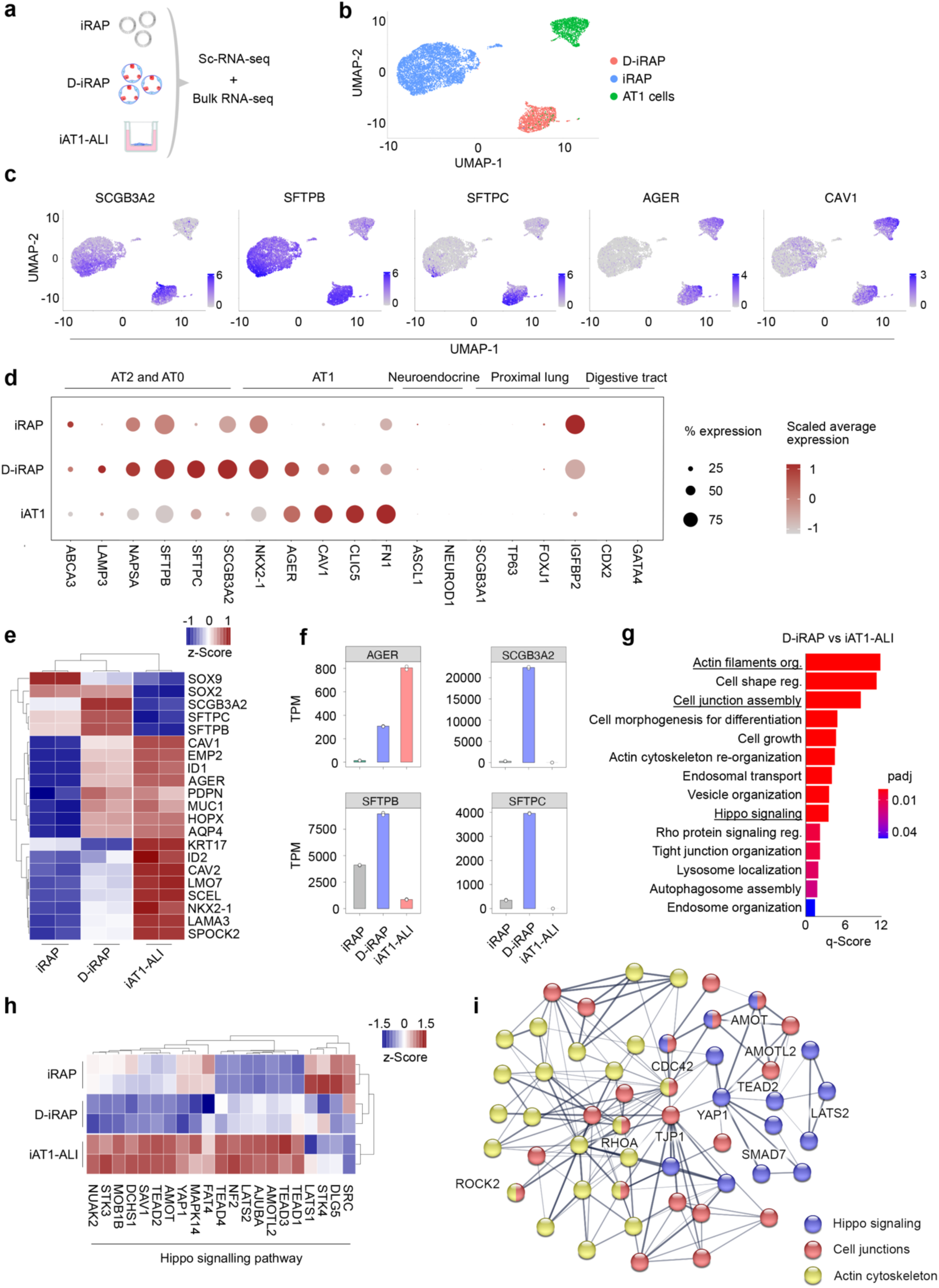
Transcriptomic analysis of iAT1 generation. **a.** Schematic of experimental design. **b.** UMAP of merged analysis of scRNA-seq of iRAPs, D-iRAPs and iAT1 cells. **c.** UMAP feature plots of indicated markers. **d.** Dot plot showing expression of indicated markers in scRNAseq of of iRAPs, D-iRAPs and iAT1 cells. **e.** Heat map of differentially expressed distal lung-associated genes from RNAseq of iRAPs, D-iRAPs and iAT1 cells. Data represent normalized counts. **f.** Expression of the indicated genes in RNAseq of iRAPs, D-iRAPs and iAT1 cells. **g.** AT1-related differentially enriched GO pathways. **h.** Heat map of expression of genes associated with Hippo signaling. Data represent normalized counts. **i.** Interaction map of differentially expressed genes belonging to actomyosin, Hippo and tight junction assembly GO pathways generated using STRING-DB.

### iRAPs with mutation associated with IPF display hallmarks of IPF

Although injury to AT2 cells is considered critical to IPF pathogenesis,^41^ we examined whether iRAPs with a mutation associated with IPF would show intrinsic abnormalities, as they share the presence of LBs and expression of SFTPB with AT2 cells. We generated iRAPs from ESCs with targeted deletion of Hermansky-Pudlak Syndrome (HPS)1. HPS is caused by abnormal trafficking of lysosome-related organelles (LROs) due to mutation in one of 11 HPS genes and share pigmentation abnormalities and platelet dysfunction, as both melanosomes and platelet delta granules are LROs.^42^ Some HPS mutations (HPS1, 2 and 4) are fully penetrant for HPS-associated interstitial pneumonia (HPSIP), an entity considered clinically indistinguishable from IPF.^43^ IPF in patients with HPS is likely caused by abnormal biogenesis and trafficking of LBs, which are also LROs. We showed previously that mutation of *HPS1* induced extensive fibrosis in hPSC-derived LOs.^16,44^

scRNA-seq of WT and *HSP1^-/-^* iRAPs (**Fig. 6a**) showed similar presence of *NKX2.1* and *SFTPB* with a small fraction of *SFTPC^+^* cells in both genotypes (**Fig. 6b**). Similar expression of distal lung markers was also confirmed by RT-qPCR **(ED9a)**. *HPS1^-/-^* iRAPs, however, contained more cells expressing *KRT17*, *COL1A1* and *COL3A1* (**Fig. 6b**), which was confirmed by RT-qPCR (**Fig. 6c**), a phenotype consistent with aBC-like cells. The doubling time of *HPS1^-/-^* iRAPs was shorter compared to parental WT iRAPs in later passages, a finding potentially consistent with their accumulation in IPF **(ED9b)**. Lysotracker Red fluorescence was reduced in *HPS1^-/-^*compared to WT iRAPs **(Fig. 6d, ED9c)**, indicative of dysfunction of the lysosomal/endolysosomal compartment (which includes LBs). Given the potential role of ER stress in IPF,^41^ we assessed the unfolded protein response of the ER (UPR^ER^). Addition of tunicamycin, an inhibitor of glycosylation that induces ER stress, increased *XBP1* splicing (**Fig. 6e**), BIP expression (**Fig. 6f**), and apoptosis **(Fig. 6g, ED9d)** in *HPS1^-/-^* compared to WT iRAPs, indicative of increased ER stress susceptibility.

**Figure 6.**
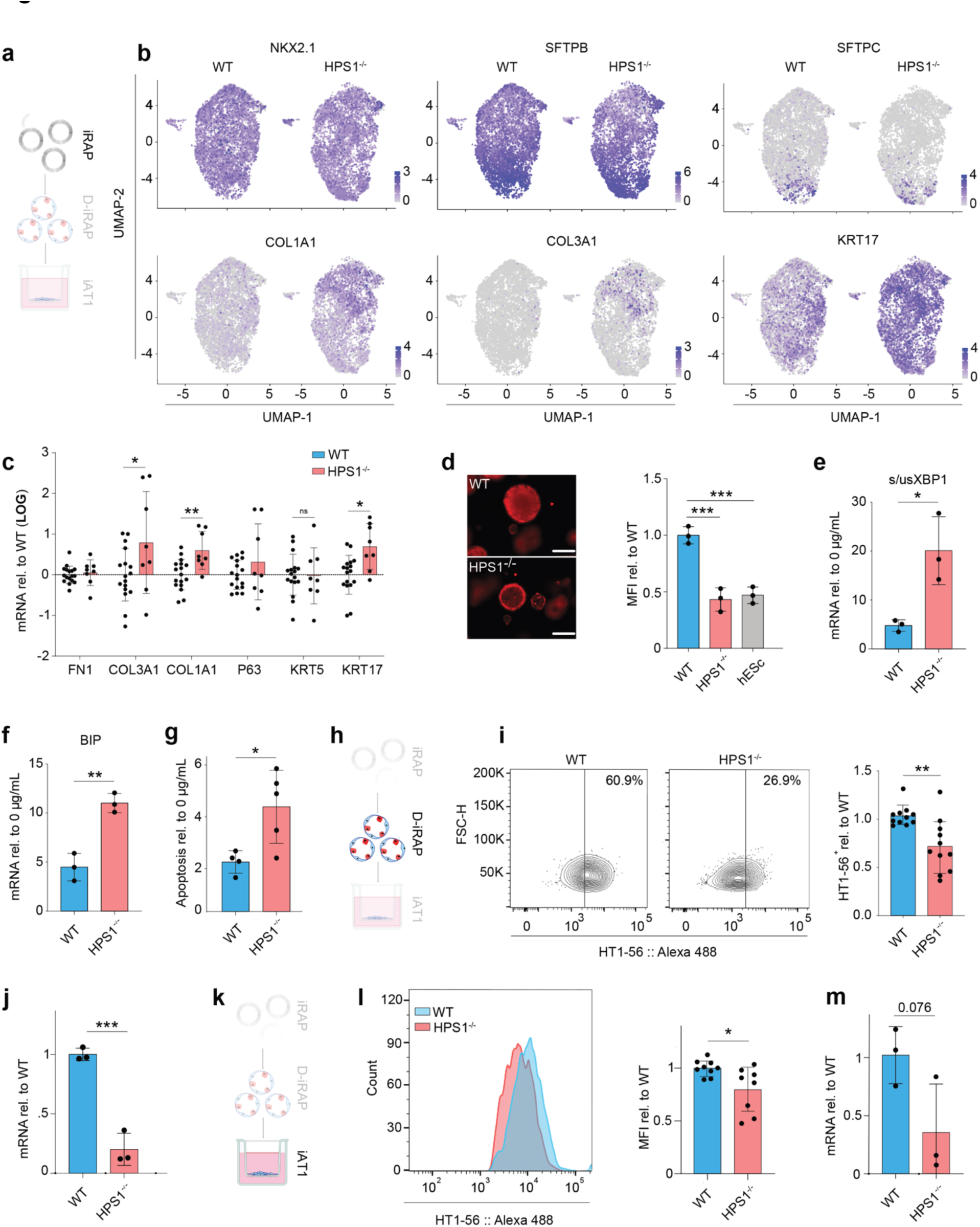
Phenotypes of *HSP1^-/-^*iRAPs. **a.** Differentiation schematic (iRAP highlighted). **b.** UMAP feature plots of indicated markers in *WT* and *HPS1^-/-^* iRAPs **c.** mRNA expression for indicated markers in *WT* and *HPS1*^-/-^ iRAPs. *p<0.05, **p<0.01, Student’s unpaired t-test, *WT* n=6, *HPS1*^-/-^ n=3. **d.** Representative images of Lysotracker^®^ uptake in iRAPs (left Scale bar = 250 µm), with flow cytometric quantification (right). ***p<0.001, One-way ANOVA with Dunnett’s multiple comparison test, n=3. **e,f.** mRNA expression for indicated markers after 5 hours of 5 ng/mL tunicamycin exposure. *p<0.05, **p<0.01, Student’s unpaired t-test, n=3. **g.** Apoptosis after 5 days of 0.1 mg/mL tunicamycin treatment normalized to untreated iRAPs of the same genotype. *p<0.05, Student’s unpaired t-test, n≥4. **h**. Differentiation schematic (D-iRAP highlighted) **i.** Representative flow cytometry plot for HT1-56 (left) with quantification of HT1-56^+^ cells in *HSP1^-/-^*D-iRAPs normalized to *WT* (right). **p<.01, Student’s unpaired t-test, n=11. **j.** mRNA expression for *AGER* mRNA in *HPS1*^-/-^ and *WT* D-iRAPs. **p<0.01, Student’s t-test, n=3. **k.** Differentiation schematic (iAT1 highlighted) **l.** Representative flow cytometry histogram of HT1-56 staining intensity (left) with quantification (right) in *WT* and *HSP1^-/-^* iAT1 cells. *p<0.05, Student’s unpaired t-test, n=9. **m.** mRNA expression for *AGER* mRNA in *HPS1*^-/-^ and *WT* iAT1 cells. Student’s t-test, n=3.

At the D-iRAP stage (**Fig. 6h**), differentiation of *HPS1^-/-^* cells was impaired especially into the AT1 lineage as evidenced by a reduced fraction of HT1-56^+^ cells (**Fig. 6i**) and a sharp reduction in *AGER* expression (**Fig. 6j**). In the AT2 lineage, a decrease in HT2-280^+^ cells was noted **(ED9e),** but expression of other AT2 markers, *LAMP3*, *SFTPB* and *SFTPC,* was not significantly affected **(ED9f)**. At the iAT1 stage (**Fig. 6k**), *HPS1^-/-^* cells, while uniformly expressing HT1-56 **(ED9g,h)**, showed a decrease in HT1-56 mean fluorescence intensity (**Fig. 6l**) and a striking reduction in *AGER* mRNA (**Fig. 6m**) compared to WT, indicative of a differentiation or maturation defect.

We conclude that *HPS1^-/-^*iRAPs display phenotypes corresponding to abnormalities observed in IPF, suggesting that intrinsic defects in RAPs play a role in the pathogenesis.

## DISCUSSION

The iRAPs described here consist of populations identified in human terminal airways, and also include cells classified as aBCs, which are mainly, but not exclusively observed in IPF.^12,15^ iRAPs can be differentiated into AT1 cells in defined conditions, and show abnormalities that are hallmarks of IPF when mutant for HPS1, thus providing important novel insights into pathogenesis of fibrotic lung disease.

Published data on the lineage relations among RAPs are conflicting. scVelo analysis of iRAPs suggests that distal BCs are the progenitors and AT0 cells the most mature population, a differentiation trajectory that aligns with the reported proximodistal location of the corresponding populations *in vivo* (**Fig. 1a**).^14^ Basil et al. identified, using a reporter gene, SCGB3A2*^+^* cells in airway organoids that lost SCGB3A2 and acquired AT2 markers when cultured in CK-DCI.^13^ In our hands, however, these conditions maintained SCGB3A2 expression and prevented AT1 and AT2 maturation. The reasons for this discrepancy are unclear. Murthy et al., on the other hand suggested, based on trajectory analysis and studies in organoids generated from adult lungs, that AT0 cells are intermediates between AT2 and AT1 cells and can also give rise to TRB-SCs. Trajectory analysis by Habermann et al.^12^ suggested that *SCGB3A2^+^* cells can be upstream of transitional AT2 cells expressing AT1 and AT2 markers, which are precursors of AT1 and AT2 cells, a hierarchy most compatible with our observations. It is likely that, given the well-established lineage plasticity occurring during lung regeneration,^1,12^ multiple distal cell types can give rise to AT1 cells during injury repair through context-dependent differentiation trajectories.

In addition to representing a model to study human distal lung regeneration, iRAPs also allow dissection of AT1 differentiation in defined conditions. This notion is supported by network analysis of DEGs during iAT1 induction, which revealed coordinated regulation of the actomyosin cytoskeleton and Hippo signaling. CDC42, a key regulator of the actomyosin cytoskeleton,^40^ induces and maintains AT1 identity in response to mechanical stress.^5,31^ Hippo signaling is involved in AT1 differentiation after various forms of injury including bleomycin, LPS, bacterial infection and unilateral pneumonectomy,^30,36,36,37^ and its downstream transcriptional regulators, YAP and TAZ, are regulated by mechanical stress and RhoGTPases.^36,37^ It is interesting to note that pathways involved in translation were downregulated during iAT1 generation **(ED8d)**, suggesting a role for translational regulation in AT1 differentiation that was not recognized before.

iRAPs with deletion of HPS1 display functional defects, including hyperresponsiveness to ER stress, increased expansion and impaired differentiation with increased presence of aBC-like cells. These findings align with results of scRNAseq studies of IPF lungs showing depletion of AT1 and AT2 cells and accumulation of aBCs,^12,15^ and suggest that, although susceptibility to AT2 injury is a factor in IPF pathogenesis,^41^ intrinsically aberrant injury repair by the distal lung progenitor compartment may play a major role as well. In particular the AT1 differentiation defect coupled with increased aBC-like cells observed in iRAPs may be important, as regions in IPF lungs with accumulated aBC-like cells lacked RAGE^+^ AT1 cells.^3^ Furthermore, a reduction of soluble RAGE in the serum is a biomarker for IPF, potentially because of the striking depletion of AT1 cells.^45^ A role for AT1 depletion in the pathogenesis of IPF is also suggested by the observation that impairment of AT1 differentiation by deletion of CDC42 after unilateral pneumonectomy in mice resulted in fibrosis with a pattern similar to that of IPF.^31^ A role for RAPs in IPF would also explain why short telomeres are the most important risk factor for IPF, as telomere dysfunction primarily affects proliferative stem cell compartments.^46^ The fact that iRAPs contain LBs and express SFTPB suggests that mechanisms involved in impaired cellular maintenance and susceptibility to injury of AT2 cells^41^ affect these regenerative populations as well.

In conclusion, the generation of iRAPs from hPSCs allowed the development of the first protocol to generate AT1 cells from hPSCs in defined conditions and to show that not only AT2 cells, but distal lung progenitors are intrinsically dysfunctional in IPF, shedding a new light on pathogenesis and providing a versatile tool for drug discovery.

## Methods

### Human lung samples

De-identified normal human lung samples were provided by the Herbert Irving Comprehensive Cancer Center Molecular Pathology Shared Resource Tissue Bank core under IRB-AAAT8682.

### Maintenance of hPSCs

Before differentiation, Rockefeller University Embryonic Stem Cell Line 2 (RUES2, passage 24-32) or mRNAi PSC lines (generated using mRNA transfection and obtained from the Mount Sinai Stem Cell Core, NY) were maintained on mouse embryonic fibroblasts (MEFs) plated at 22,500 cells/cm^2^. Cells were cultured in hESC maintenance media (DMEM/F12 (ThermoFisher, Carlsbad, CA), 20% Knock-out serum, (Stem Cell Technologies, Vancouver, BC), 0.1 mM β-mercaptoethanol (Sigma-Aldrich,Burlington, MA), 0.2% Primocin (InvivoGen, San Diego, CA), 20 ng/ml FGF2 (R&D Systems/Biotechne, Minneapolis, MN), and 1% Glutamax (ThermoFisher)) which was changed daily. hPSCs were passaged every 3-4 days with Accutase (Innovative Cell Technologies, San Diego, CA) at least two times before differentiation, washed and replated at a dilution of 1:24. Cultures were maintained in a humidified 5% CO_2_ atmosphere at 37°C. Lines are karyotyped and verified for Mycoplasma contamination using PCR (InVivoGen) every 6 months.

### Generation of hESC-derived lung organoids

Human LOs were generated as described previously. Briefly, MEFs were depleted by passaging 5-7x10^6^ hESCs onto Matrigel (Corning, Corning, NY) coated 10-cm dish. Cells were maintained in hESC media in a humidified 5% CO_2_ atmosphere at 37°C. After 24 hours, cells were detached with 0.05% Trypsin/EDTA (ThermoFisher) and distributed to 6-well low-attachment plates containing primitive streak/embryoid body media (10 μM Y-27632 (Tocris/Biotehne, Minneapolis, MN), 3 ng/ml BMP4 (R&D/Biotechne) to allow embryoid body (EB) formation. EBs were fed every day with fresh endoderm induction media (10 μM Y-27632, 0.5 ng/ml BMP4, 2.5 ng/ml FGF2 and 100 ng/ml ActivinA (R&D/Biotechne)) and maintained in a humidified 5% CO_2_/5% O_2_ atmosphere at 37°C. Endoderm yield was determined by dissociating EBs and evaluating CXCR4 and c-KIT (Biolegend, San Diego, CA) co-expression by flow cytometry on day 4. Cells used in all experiments had > 90% endoderm yield and were plated on 0.2% fibronectin-coated wells (R&D/Biotechne) at a density of 80,000 cells/cm^2^. Cells were incubated in Anteriorization media-1 (100 ng/ml Noggin (R&D/Biotechne) and 10 μM SB431542 (Tocris)) for 24 hours, followed by Anteriorization media-2 (10 μM SB431542 (Tocris) and 1 μM IWP2 (Tocris)) for another 24 hours. At the end of anterior foregut endoderm induction, cells were switched to Ventralization/Branching media (3μM CHIR99021 (Tocris), 10 ng/ml FGF10 (R&D/Biotechne), 10 ng/ml rhKGF, (R&D/Biotechne), 10 ng/ml BMP4 and 50 nM *all-trans* Retinoic acid (Tocris)) for 48 hours and three-dimensional clump formation was observed. The adherent clumps were detached by gentle pipetting and transferred to the low-attachment plate, where they folded into lung bud organoids (LBOs) at d10-d12. Branching media was changed every other day until d20–d25 and LBOs were embedded in 100% Matrigel in 24-well transwell (BDFalcon, Franklin Lakes, NJ) inserts. Branching media was added after Matrigel solidified and changed every 2-3 days to facilitate proper growth into lung organoids. Culture of embedded organoids can be kept for more than 6 months.

### Generation of induced respiratory airway progenitors (iRAP)

Matrigel embedded lung organoids can be used for iRAP generation when they reach d42 of development. Media was removed from the transwell and 1 ml of 2 mg/ml dispase (Corning) was added to release lung organoid from the Matrigel for 30-45 minutes in a normoxic incubator. The organoid was transferred to a 15 ml conical tube and washed with stop media (DMEM (Corning), 5% FBS (Atlanta Biologicals; Flowery Branch, Georgia), 1% Glutamax (ThermoFisher)) to neutralize dispase, then centrifuged at 200 g for 5 minutes. The pellet was incubated with 1 ml of 0.05% Trypsin/EDTA in a normoxic incubator for 10-12 minutes with occasional pipetting with a P1000 pipet tip. Single cell dissociation was verified using a bright field microscope. If a single cell suspension was not obtained after 12 minutes, cells were washed with stop media and incubated for additional 5 minutes with 0.05% Trypsin/EDTA. Cells were counted using a hemocytometer and 400 cells/μl of undiluted Matrigel were plated in a well of a 12-well non-tissue culture plate. The plate was placed in normoxic incubator for 30 minutes until Matrigel polymerized and 1 ml of CK-DCI (3μM CHIR 99021, 10 ng/ml rhKGF, 50 ng/ml dexamethasone (ThermoFisher), 0.1 mM 8-Bromoadenosine 3′,5′-cyclic monophosphate sodium salt (Tocris) and 0.1 mM 3-Isobutyl-1-methylxanthine (Sigma-Aldrich)) media was gently added using 10 ml serological pipette. Media was changed every 3 days. After 2-3 weeks, an iRAP culture is established that can be maintained for at least 8 passages.

### Generation of D-iRAPs

7-10 days after passaging, small iRAP spheres formed. CK-DCI media was replaced with DCI-SB ((50 ng/ml dexamethasone (ThermoFisher), 0.1 mM 8-Bromoadenosine 3′,5′-cyclic monophosphate sodium salt (Tocris) and 0.1 mM 3-Isobutyl-1-methylxanthine (Sigma-Aldrich), 10 µM SB-431542 (Tocris)) for 10-14 days.

### Generation of iAT1 cells

6-well transwell inserts (BDFalcon, Franklin Lakes, NJ) were coated with 1:15 Matrigel:SB-BMP4-RI (10 µM SB-431542 (Tocris), 10 ng/mL BMP4 (Tocris), and 10 µM Y-27632 (Tocris)) and left at 4°C overnight. Mature D-iRAPs were released from Matrigel using dispase and washed, as described above. Media was aspirated from the transwell when ready to seed with cells. 2.5 x 10^5^ cells were resuspended in 1 mL of SB-BMP4-RI and added into the insert. 1-2 mL of SB-BMP4-RI was added into the 6 well. The cells were cultured for 5-7 days in a normoxic incubator at 37°C. For ALI, 7 days after seeding on the transwell insert, the media in the insert was removed and the cells were incubated for another 4-5 days.

### RT-qPCR

Total RNA was extracted according to manufacturer’s instructions using the Direct-zol™ RNA Microprep (Zymo Research, Irvine, CA) and 500 ng of total RNA was reverse transcribed using the qScript™ XLT cDNA SuperMix (Quantabio, Beverly, MA). Technical triplicates of 15 μl reaction (for use in Applied Biosystems QuantStudio7 384-well System, Waltham, MA) were prepared with 3 µl of diluted cDNA and run for 40 cycles. Primers used are listed in **Table S1**. Relative gene expression was calculated based on the average cycle (Ct) value, normalized to *GAPDH* as the internal control and reported as fold change (2(-ddCT)).

### Immunofluorescence

After removal of media, a cell scraper was used to carefully detach the Matrigel droplet with embedded IRAPs from the bottom of the 12 well plate. The droplet was then transferred to an OCT histology mold and embedded in Tissue-Tek OCT (Sakura Finetek, Torrance, CA). Frozen samples were cut using a cryotome at the thickness of 10-12 μm and collected on adhesion microscope slides, air-dried and fixed in 4% paraformaldehyde, then washed 2 times for 5 minutes in 50 mM glycine to inactivate PFA, followed by washing in PBS. Samples were permeabilized for 10 minutes in 0.2% PBST (PBS + 0.2% Triton X-100 (Fisher Scientific, Hampton, NH)) and blocked by incubating in PBS containing 5% donkey serum, then incubated overnight in primary antibody **(Table S2)** in 0.2% Triton X-100 and 2% donkey serum. On the next day, samples were washed three times in PBS and 1% donkey serum and incubated with secondary antibodies (1:200, **Table S2**) for 1 hour at room temperature. Nuclei were stained with DAPI (ThermoFisher) and sections were mounted with Mounting Reagent (DAKO, Santa Clara, CA) on coverslips. Samples were imaged using a Leica TCS SP8 Stellaris Laser scanning confocal microscope, and Leica DMi1 Inverted Phase Contrast Microscope (Leica Microsystems, Deerfield, IL).

### 3D imaging

Tips and conical 15 ml plastic tubes were pre-coated with 1% (wt/vol) PBS–BSA. D-iRAPs were released from Matrigel as previously described in the ‘Generation of induced Respiratory Airway Progenitor’ section. Spheres were transferred into a precoated 15 ml falcon tube with ice cold 1% (wt/vol) PBS–BSA. Tubes were centrifuged at 70g for 3 min at 4°C. Pellets were resuspended in ice cold 4% PFA with precoated tips and incubated for 10 min. 15 ml conical plastic tubes were filled with PBT (PBs-Tween: 0.1% (wt/vol) Tween 20 in PBS) for 10 min at 4°C on roller rotation. Spheres were centrifuged at 70g for 3 min at 4°C and resuspended with ice cold 1% (wt/vol) PBS– BSA for 45 min at 4°C on roller rotation. Staining procedure followed the same step as described in the ‘Immunofluorescence’ section, albeit both primary and secondary antibodies were incubated overnight at 4°C on roller rotation. Mounting was performed on a glass slide, between two coverslips. Imaging was performed on a Leica TCS SP8 Stellaris Laser scanning confocal microscope. Z-stacks were acquired every ∼700 nm, covering the height of each sphere.

### Image processing

The 3D reconstituted images were analyzed using Aivia software version 12.0.0. The intensity detection threshold for each signal was set using iRAPs signal intensity. Surfaces were created for DAPI, AGER, proSFTPC and SFTPC signals. Surface parameters consisted of an image smoothing filter size of 9, a minimum edge intensity of 30 and a partition radius of 20 μm. Surface measurements were collected and normalized to control conditions. 2D images were processed using ImageJ version 2.1.0/1.53c.

### ER stress

CK-DCI media was replaced with CK-DCI plus 5 µg/mL tunicamycin (Sigma-Aldrich) and returned to a normoxic incubator for 5 hours. Cells were then collected for mRNA analysis. For apoptosis studies, CK-DCI media was replaced with CK-DCI plus 0.1 µg/mL tunicamycin (Sigma-Aldrich) and returned to a normoxic incubator for 5 days. Cells were then released and prepared for cell-cycle analysis (see below).

### Lysotracker uptake

CK-DCI media was replaced with CK-DCI plus Lysotracker Red DND-99 (Thermofisher) at a concentration of 75 nM and returned to a normoxic incubator for 2.5 hours. Cells were kept in Matrigel and imaged using Leica DMi1 Inverted Phase Contrast Microscope (Leica Microsystems, Deerfield, IL) and then released from Matrigel for flow cytometric analysis.

### Flow cytometry

Spheres embedded in Matrigel were released by incubating with dispase for 30-60 minutes, then washed and dissociated into single cells with 0.05% Trypsin/EDTA in a normoxic incubator for 10-12 minutes with occasional pipetting with a P1000 tip. The single cell suspension was stained in polystyrene round-bottom 12 x 75 mm tubes (BD Falcon). Primary HT1-56 (Terrace Biotech, San Francisco, CA, 1:200 dilution) and HTII-280 (Terrace Biotech, San Francisco, CA, 1:200 dilution) antibodies were incubated for 1 hour at room temperature. Cells were washed two times with FACS buffer (PBS, 1% BSA) and centrifuged for 5 minutes at 1400 rpm. Fluorochrome-labeled secondary antibody (Alexa Fluor 488 #A21042, or 594 #A21044 goat anti-mouse IgM, Thermofisher) were diluted in FACS buffer at 1:100 and added for 30 minutes in the dark. Cells are washed twice. Conjugated human EPCAM antibody (Biolegend) was added for 30 minutes. Cells were washed and resuspended in FACS buffer for flow cytometric analysis on a ZE5 cell analyzer (Bio-Rad, Hercules, CA) or a Novocyte Quanteon (Agilent, Santa Clara, CA).

### Cell Cycle Analysis

Single-cell suspensions were prepared as described above. Single cells were resuspended in 200 µL PBS and ethanol-fixed by adding 800 µl of ice-cold 100% ethanol drop-wise while gently vortexing. Cells were left in fixative for at least 2 hours. Fixed cells were washed in 2 mL PBS 2x after which 200 µL of freshly made propidium iodide solution (.1% Triton-X 100 (Fisher Scientific), 50 µg/mL propidium iodide (Thermo Scientific), 0.2 mg/mL RNAse A (Thermo Scientific) in PBS) was added. Samples were incubated at 37°C for 20 min before flow cytometric analysis.

### Transmission Electron Microscopy

Transmission Electron Microscopy (TEM) was performed at the NYU Langone Medical Center Microscopy Core. iRAPs were fixed with 2% paraformaldehyde and 2.5% glutaraldehyde in 0.1M sodium cacodylate buffer (pH 7.2) for 2 hours and post-fixed with 1% osmium tetroxide for 1.5 hours at room temperature, then processed in a standard manner and embedded in EMbed 812 (Electron Microscopy Sciences, Hatfield, PA). Semi-thin sections were cut at 1 μm and stained with 1% Toluidine Blue to evaluate the quality of preservation and find the area of interest. Ultrathin sections (60 nm) were cut, mounted on copper grids and stained with uranyl acetate and lead citrate by standard methods. Stained grids were examined under Philips CM-12 electron microscope and photographed with a Gatan (4k ×2.7k) digital camera (Gatan, Inc., Pleasanton, CA).

### Western Blot

Cells embedded in Matrigel were released by incubating with dispase (Corning) for 30 minutes. Cells were washed, collected in PBS, and lysed in lysis buffer (RIPA buffer, 1x Roche Complete Protease inhibitor cocktail, and Roche PhosSTOP). Buffer-treated cells were mechanically lysed using a 25-gauge needle and left on ice for 30 minutes. Human lung samples were homogenized with stainless steel beads and left on ice in the lysis buffer for 4-6 hours. Samples were centrifuged at 15,000 g for 10 minutes and the supernatant was collected and stored at -80°C until analysis. Protein concentration was measured using the Pierce BCA Protein Assay Kit (ThermoFisher). A total of 18 μg of iRAP, hPSC, and human lung lysate were loaded for mature SFTPC analysis and a total of 5 μg of each were loaded for mature SFTPB analysis. Lysates were resolved on pre-cast NuPage 4-12% Bis-Tris gels (Invitrogen) and transferred to nitrocellulose membranes (Bio-Rad, Hercules, CA). The following primary antibodies were used to probe the blots: mature SFTPC (1:1000, Seven Hills Bioreagents, Cincinatti, OH), mature SFTPB (1:1000, Seven Hills Bioreagents), GAPDH (1:5000, Cell Signaling Technology). Species-specific secondary antibodies with HRP conjugates (Cell Signaling Technology, Danvers, MA) were used at 1:15,000 dilution. Blots were n treated with Pierce ECL Western Blotting Substrate and visualized using the Amersham ImageQuant 800 (Cytiva, Marlborough, MA).

### Cryopreservation of iRAPs

2-3 weeks after iRAPs were passaged, CK-DCI media was removed and 1 mL of dispase (2 mg/mL) released the spheres. After 30-45 minutes in a normoxic incubator, the spheres were transferred to a 15 mL conical and washed with wash media, then centrifuged at 200 g for 5 minutes. Wash media was aspirated and the pellet was incubated with 1 mL of 0.05%Trypsin/EDTA in a normoxic incubator for 10-12 minutes. They were washed again, followed by resuspension of the pellet in a small volume of CK-DCI media. Cells were frozen at a density of 5x10^5^ to 10^6^ single cells. Cells are resuspended in equal volumes of CK-DCI media and 2x DMSO freezing medium (Quality Biological, Gaitersburg, MD), transferred to a cryovial, and placed in a container that allows a freezing rate of -1°C/min in a -80°C freezer. The next day, cells were transferred to liquid nitrogen. When ready to use, the cryovial is placed in a 37°C water bath until thawed. Cells are transferred to a 15 mL conical tube and washed with wash media, centrifuged at 200 g for 5 minutes, and aspirated. The ideal initial reseeding density is 800 - 1,600 cells per μL of undiluted Matrigel.

### RNA-seq and data analysis

Total RNA was extracted from iRAPs, D-iRAPs and iAT1 samples using Qiagen RNA extraction kit according to the manufacturer’s instructions. RNA quality was evaluated by Agilent 2100 BioAnalyzer. RNA-seq libraries were prepared and sequenced by the JP Sulzberger Columbia Genome Center High-Throughput Screening Center at Columbia University. Paired-end equencing was performed on a NovaSeq 6000 platform. RNA-seq read quality assessment was performed using fastQC v.0.12.0. Processed reads were mapped against the human genome (GRCh38.p14) using Gencode v44. The number of reads per gene was counted using Salmon v.1.10.12 providing the count matrix. Differential gene expression was analyzed using DESeq2 v.1.30.1 R package Differentially expressed genes were identified for our study by setting a cutoff of pValue < 1*10^-10 and a fold-change > ±2. Gene ontology analysis was performed using the differentially expressed genes with enrichplot package v.1.10.2, using genome-wide annotation for human database v.3.12.0. Genes contributing to GO terms were fed into String-DB software to perform the protein network analysis according to the pipeline described in ED7e.

### Single-cell cDNA library preparation and scRNA-seq

Samples were released from Matrigel using dispase and dissociated into single cells using trypsin. For the multiplexing experiment, dissociated D-iRAPS and iAT1 were tagged using TotalSeq-B anti-human hashtags. The viability of single cells was assessed using Trypan Blue staining, and debris-free suspensions of >80% viability were deemed suitable for single-cell RNA Seq. sc-RNA-seq was performed using the Chromium platform (10x Genomics, Pleasanton, CA) with the 3’ gene expression (3’ GEX) V3 kit, using an input of ∼10,000 cells. Briefly, Gel-Bead in Emulsions (GEMs) were generated on the sample chip in the Chromium controller. Barcoded cDNA was extracted from the GEMs by Post-GEM RT-cleanup and amplified for 12 cycles. Amplified cDNA was fragmented and subjected to end-repair, poly A-tailing, adapter ligation, and 10X-specific sample indexing following the manufacturer’s protocol. Libraries were quantified using Bioanalyzer (Agilent) and QuBit (Thermofisher) analysis and were sequenced in paired end mode on a NovaSeq instrument (Illumina, San Diego, CA) targeting a depth of 50,000-100,000 reads per cell.

### scRNA-seq computational analysis

Sequencing data were aligned and quantified using the Cell Ranger Single-Cell Software Suite (version 6.1.2, 10x Genomics) against the provided GRCh38 (Ensembl 98) human reference genome. All further computational analysis of scRNA-seq data was performed using R version 4.1.3 (https://www.R-project.org/.) unless otherwise stated.

#### QC and processing

The aligned data was imported and processed using the R package Seurat v4.1.1^38^. Quality control for doublets and low-quality cells was achieved through exclusion of cells with less than 500 or more than 9000 transcripts and those with a higher than 20% mitochondrial gene contribution, respectively. Additionally, transcripts were retained if they counted over 0 in more than 0.5 % of all cells, otherwise excluded. Count data was then log-normalized and transcripts were scaled and centered, using built-in Seurat functions. Variable transcripts were calculated based on standardized feature values using observed mean and expected variance of a local polynomial regression model. On the resulting variable transcripts 50 principal components were computed, which in turn were used as input for uniform manifold approximation and projection (UMAP) dimensionality reduction. For clustering analysis, a shared nearest neighbor (SNN) graph was constructed and the modularity function optimized using the Leiden algorithm.

#### Cell type annotation

Potential cell types present in our dataset were predicted through machine learning. In brief, a random forest classifier (SingleCellNet R package)^47^ was trained on fully annotated published data by Murthy et al.^14^ and Haberman et al.^12^ and assessed on a withheld subset. This classifier was then applied to the current dataset and a matching cell type was predicted for each cell. Azimuth-based cell type identification using Human Lung Cell Atlas^34^ was used for iAT1 samples.

#### RNA velocity

RNA velocity was calculated in python using the packages Velocyto and scVelo, according to the developer’s manual Briefly, ‘.loom’ files containing both exon and intron information were created from aligned raw data using Velocyto. scVelo was then used to normalize the data and compute moments. Subsequently, RNA velocity was estimated and projected onto Seurat-derived UMAP coordinates. The length and coherence of the velocity vectors, which indicate differentiation speed and directional confidence, respectively, were calculated. Finally, a dynamical model was applied to analyze transcriptional states and cell-internal latent time and subsequently recompute RNA velocities. The latent time, based solely on the transcriptional dynamics of a cell, was thereby determined.

### Statistical analysis

Statistical analysis was performed using unpaired two-tailed Student’s t-test, one-way ANOVA, or two way-ANOVA where appropriate using Prism 9. Results are shown mean ± SD. P-values < 0.05 were considered statistically significant. N-value refers to biologically independent replicates, unless noted otherwise.

## Supporting information

Source Data: Differentially expressed genes during AT1 differentiation

## Acknowledgments

This work was supported by grants NIH HL120046 (HWS), NIH 1U01HL134760 (HWS), NIH S10 OD032447 (HWS), and NIH T32GM145440 (JAT). This research was also funded in part through the NIH/NCI Cancer Center Support Grant P30CA013696 and used the Tissue bank part of the Molecular Pathology Shared Resource.

## Author Contributions

MGP and JAT have performed most of the experiments and analysis, IML developed the generation of iRAPs, JWM oversaw microscope analysis. TAT and NS performed scRNA-seq analysis. KGB and HWS supervised scRNA-seq work and analysis. HWS provided concept, supervised and wrote the manuscript.

## Data accessibility

Bulk and single cell RNAseq data have been deposited in the GEO database, accession GSE245723: Reviewer’s token atcvgyyyrbwtjsv.

---

**ED1.**
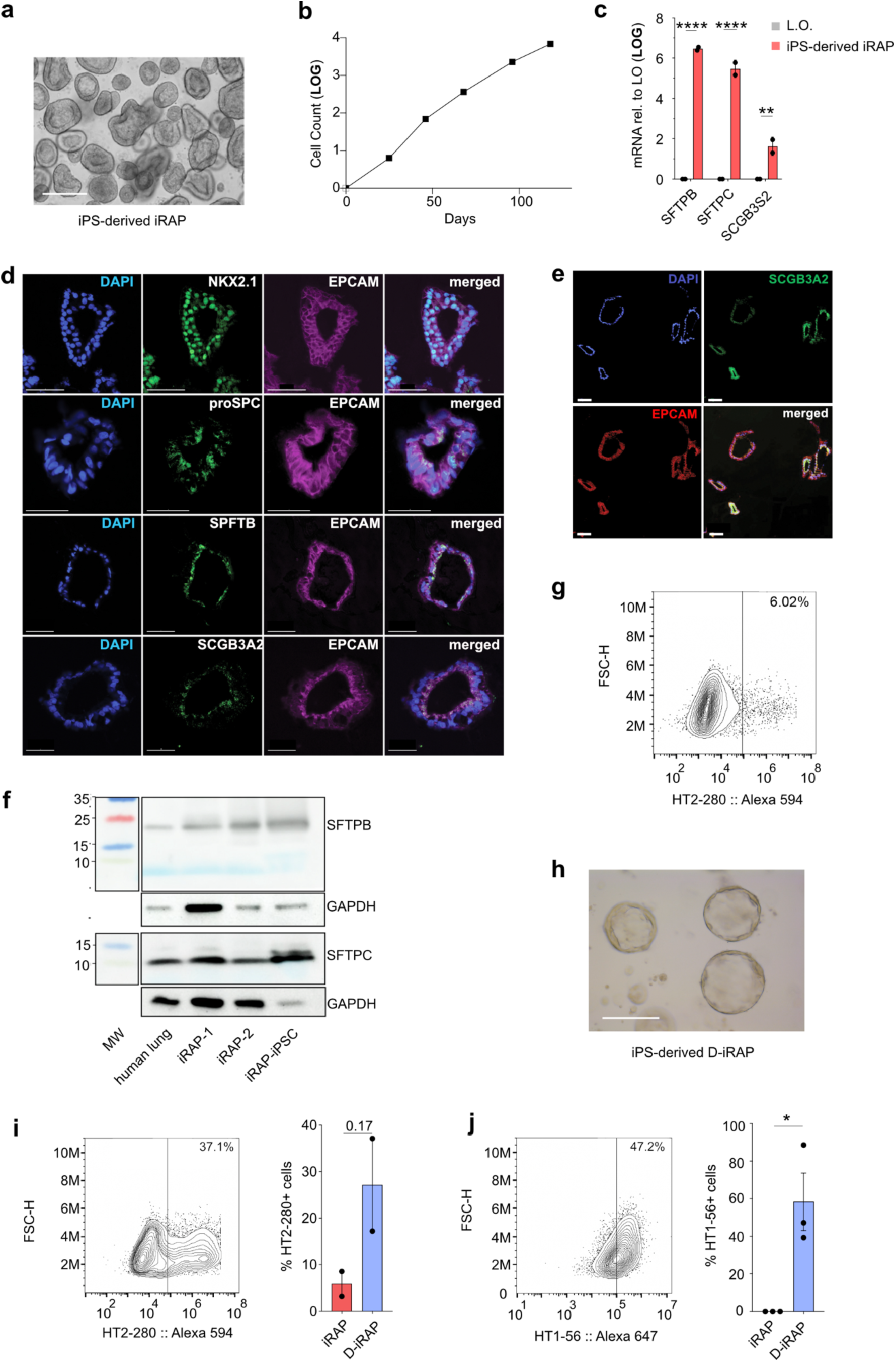
iPSC-derived iRAPs. **a.** Representative bright-field image of iPS-derived iRAPs (scale bar = 400 µm). **b.** Representative expansion of iPSC-derived iRAPs (n=1). **c.** mRNA expression for indicated markers in iPSC-derived iRAPs compared to LOs. ** p<0.01, **** p<0.0001 two-way ANOVA with Sidak test for multiple comparisons, n=2. **d-e.** Representative confocal images for indicated markers of iRAPs generated from iPSCs. Scale bar = 50 µm. **f.** Representative WB for mature SFTPB and SFTPC in iPSC- and ESC-derived iRAPs and human adult lung. **g.** Representative flow cytometry plot for HT2-280. **h.** Representative bright-field image of iPS-derived D-iRAPs. Scale bar = 200 µm. **i.** Representative flow cytometry plot of iPS-derived D-iRAPs for HT2-280 (left) with quantification (right). Student’s t-test, n=2). **j.** Representative flow cytometry plot for iPS-derived D-iRAPs HT1-56 (left) with quantification (right). *p<0.05, Student’s t-test, n=3.

**ED2.**
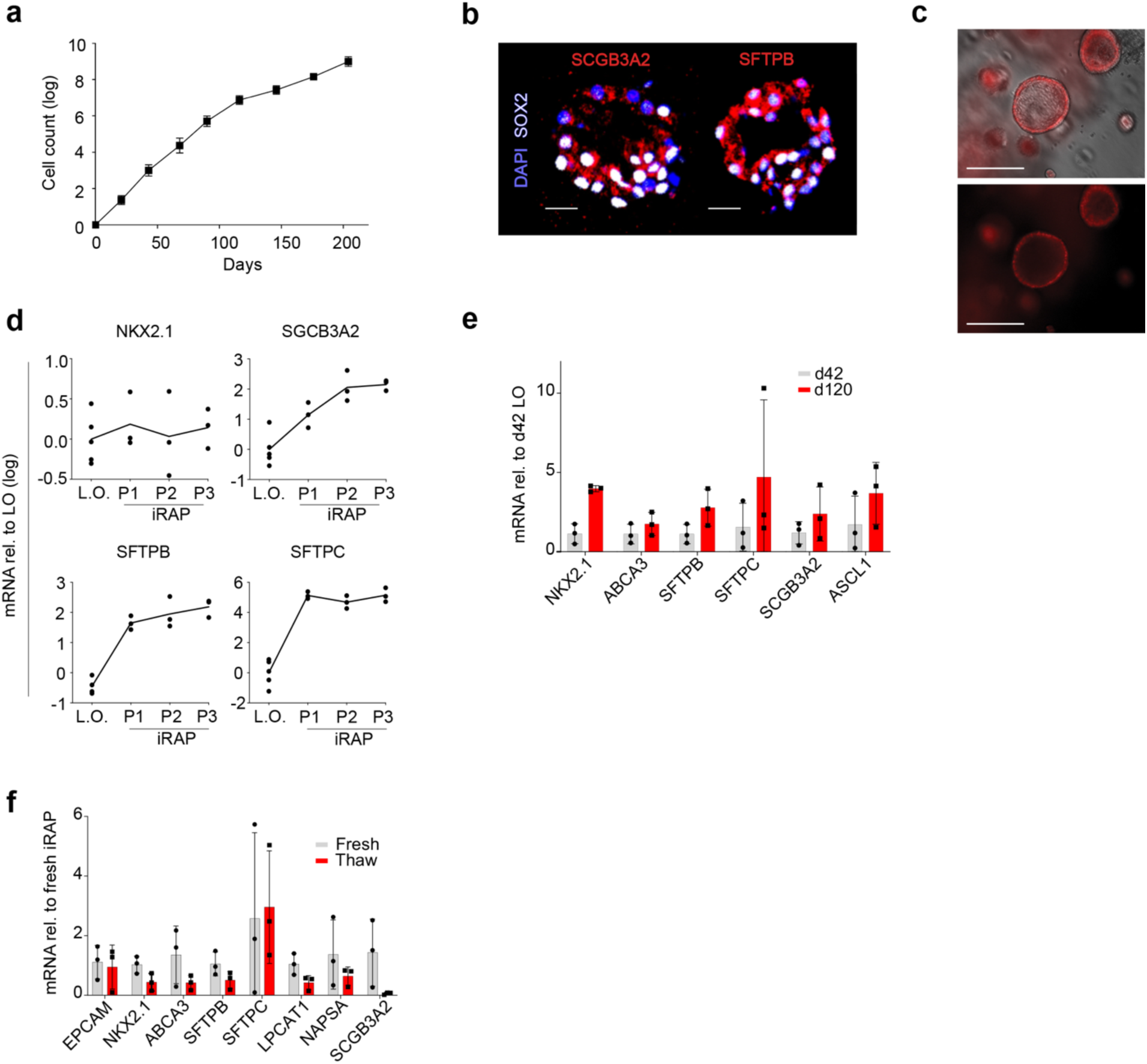
Further characterization of iRAPs. **a.** Expansion of ESC-derived iRAPs (n=3). **b.** Representative confocal image for selected markers. Scale bar = 50 µm **c.** Representative image of Lysotracker Red uptake. Scale bar = 200 µm **d.** Expression mRNA for indicated genes at consecutive passages relative to LOs. **e.** mRNA expression for lung-specific markers in ESC-derived iRAPs derived from day 120 relative to from d42 LOs. Two-way ANOVA with Sidak’s multiple comparison test (n=3). **f.** mRNA expression for lung-specific markers in ESC-derived iRAPs after cryopreservation relative to fresh iRAPs. Two-way ANOVA with Sidak’s multiple comparison test (n=3).

**ED3.**
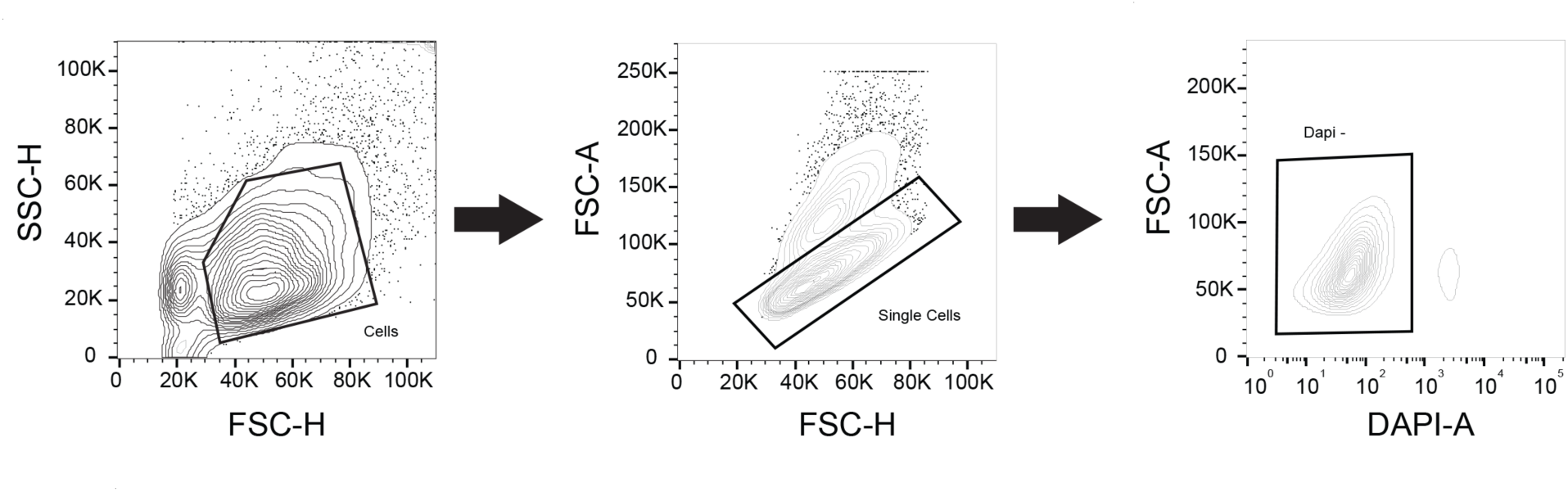
Gating strategy. Representative example of live cells gating and doublet exclusion used throughout this report.

**ED4.**
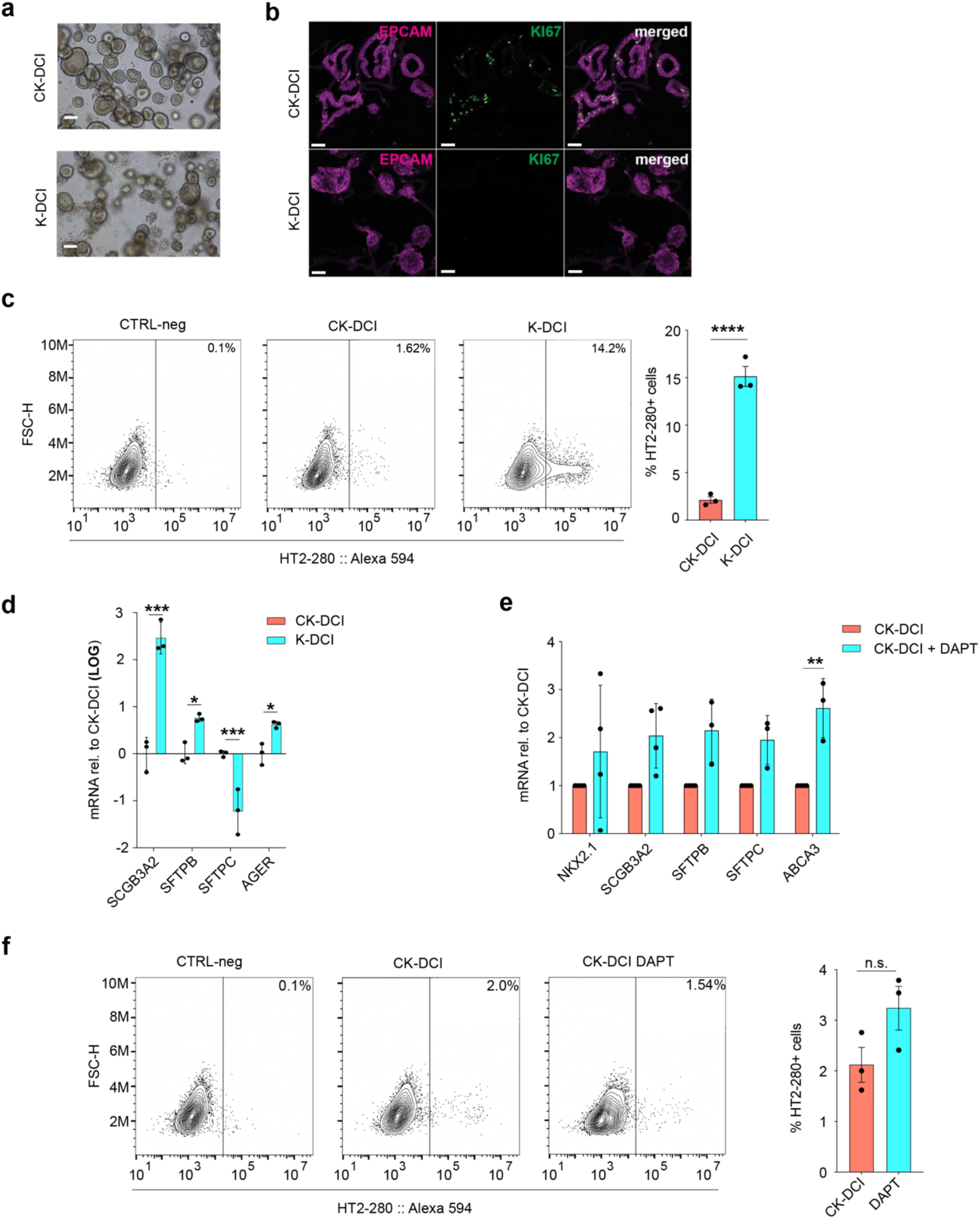
Establishment of conditions to induce D-iRAPs. **a.** Representative bright field images after CHIR withdrawal. Scale bar = 200 µm. **b.** Representative IF for indicated markers. Scale bars = 100 µm. **c.** Representative flow cytometry plot (left) with quantification of HT2-280^+^ cells (right). **** p<0.0001, Student’s t-test, n=3. **d.** mRNA expression for indicated markers after CHIR withdrawal. ***p<0.001, two-way ANOVA with Sidak multiple comparison test (n=3). **e.** RT-qPCR for after addition of DAPT. *** p<0.001, Two-way ANOVA with Sidak multiple comparison test (n=3). **f.** Representative flow cytometry plot (left) with quantification (right) of HT2-280^+^ cells. Student’s t-test, n=3.

**ED5.**
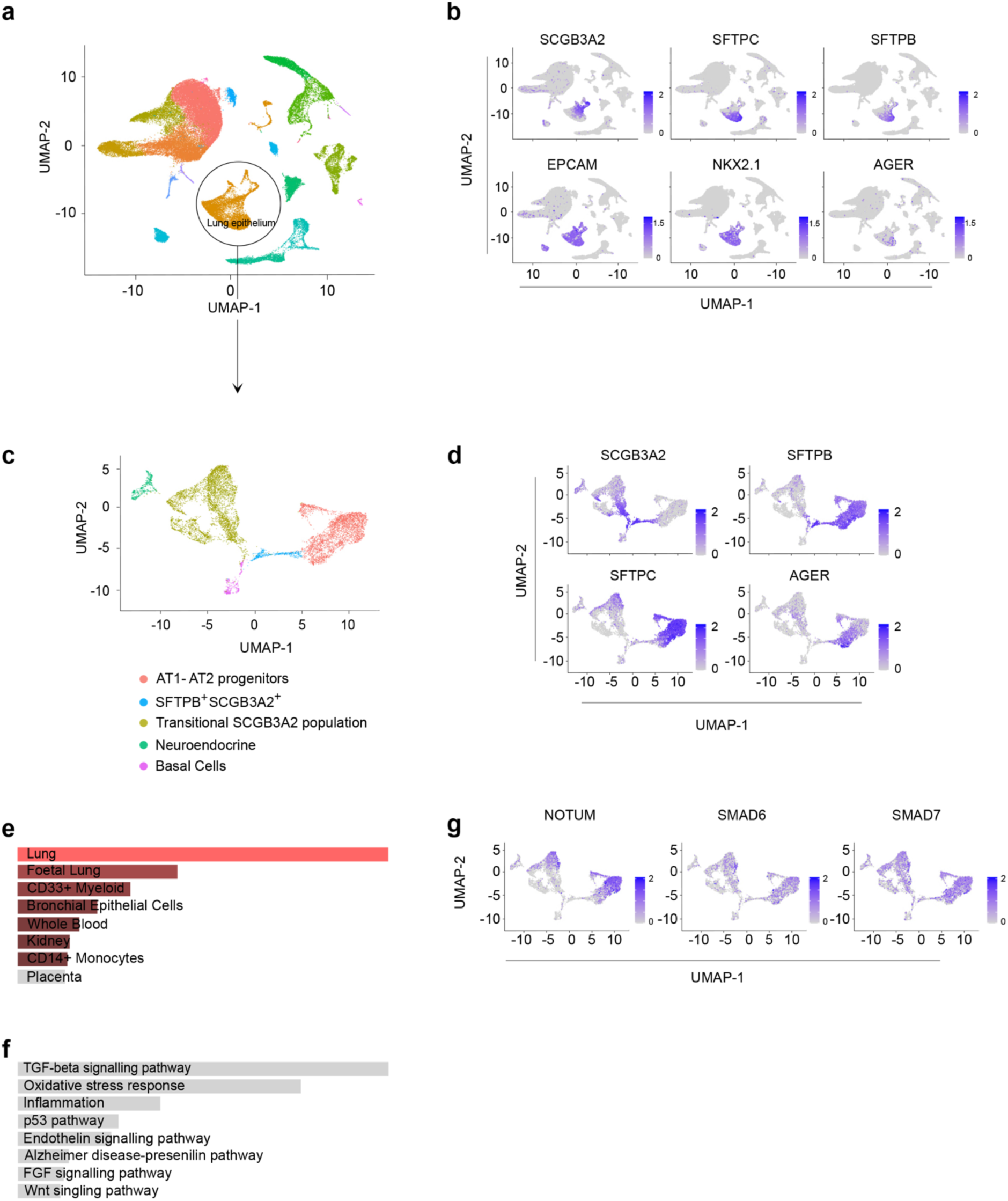
Signaling pathways involved in human AT1 and AT2 development. **a.** Cluster analysis of human embryonic lung sc-RNA-seq dataset. “Lung epithelium” cluster identity based on gene expression analysis in (**b**). **b.** UMAP feature plots for indicated markers. **c.** Cluster analysis after sub-setting ‘Lung epithelium’ cluster. Cluster identity based on gene expression in **(d)**. **d.** UMAP feature plots for indicated markers. **e.** Cell type analysis 100 top DEGs in “AT1-AT2 progenitor” cluster. **f.** GO term analysis (Panther) of 100 top DEGs in “AT1-AT2 progenitor” cluster. **g.** UMAP feature plots for indicated markers.

**ED6.**
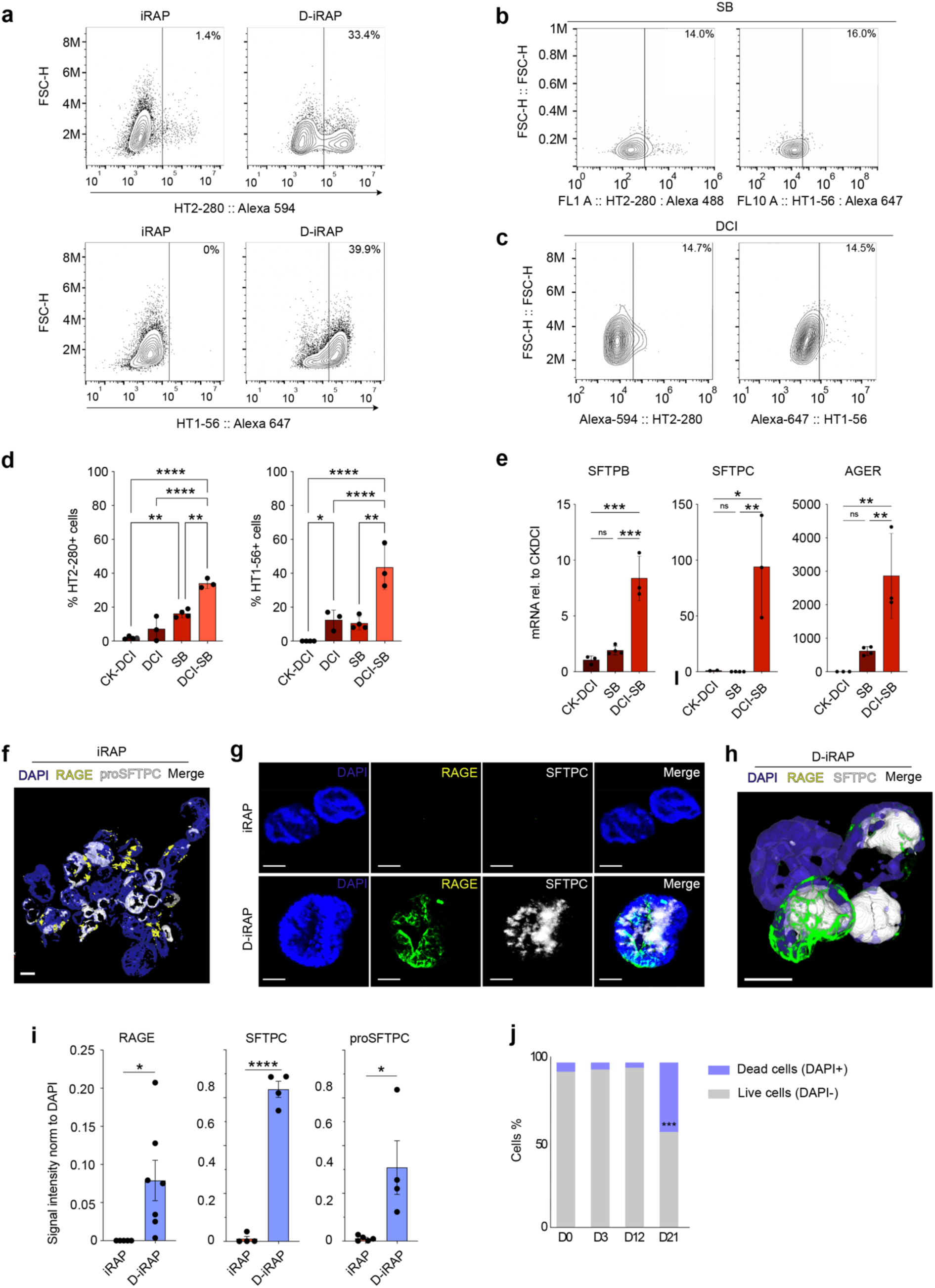
Further characterization of D-iRAPs. **a-c.** Representative flow cytometry plots for indicated markers and conditions. **d.** Quantification of flow cytometry shown in **(b)** and **(c)**. *p<0.05, **p<0.01, ****p<0.0001, one-way ANOVA with Tukey multiple comparison test (n=3). **e.** mRNA expression for indicated markers in conditions shown in X-axis. *p<0.05, **p<0.01, ***p<0.001, one-way ANOVA with Tukey multiple comparison test. **f & h.** Surface reconstitution based on indicated marker fluorescent intensity. Scale bar = 150 µm**. g.** 3D reconstitution of representative confocal 3D images for indicated markers. Scale bars = 50 µm. **i.** Fluorescent signal intensity quantification of the indicated markers. *p<0.05, ****p<0.0001, Student’s t-test, n=42 iRAPs and n=97 D-iRAP spheres from 4 independent replicates. **j.** Cell viability during differentiation of D-iRAPs from iRAPs in DCI-SB conditions for up to 21 days, ***p<0.001, two-way ANOVA with multiple comparison using Sidak test, n=3.

**ED7.**
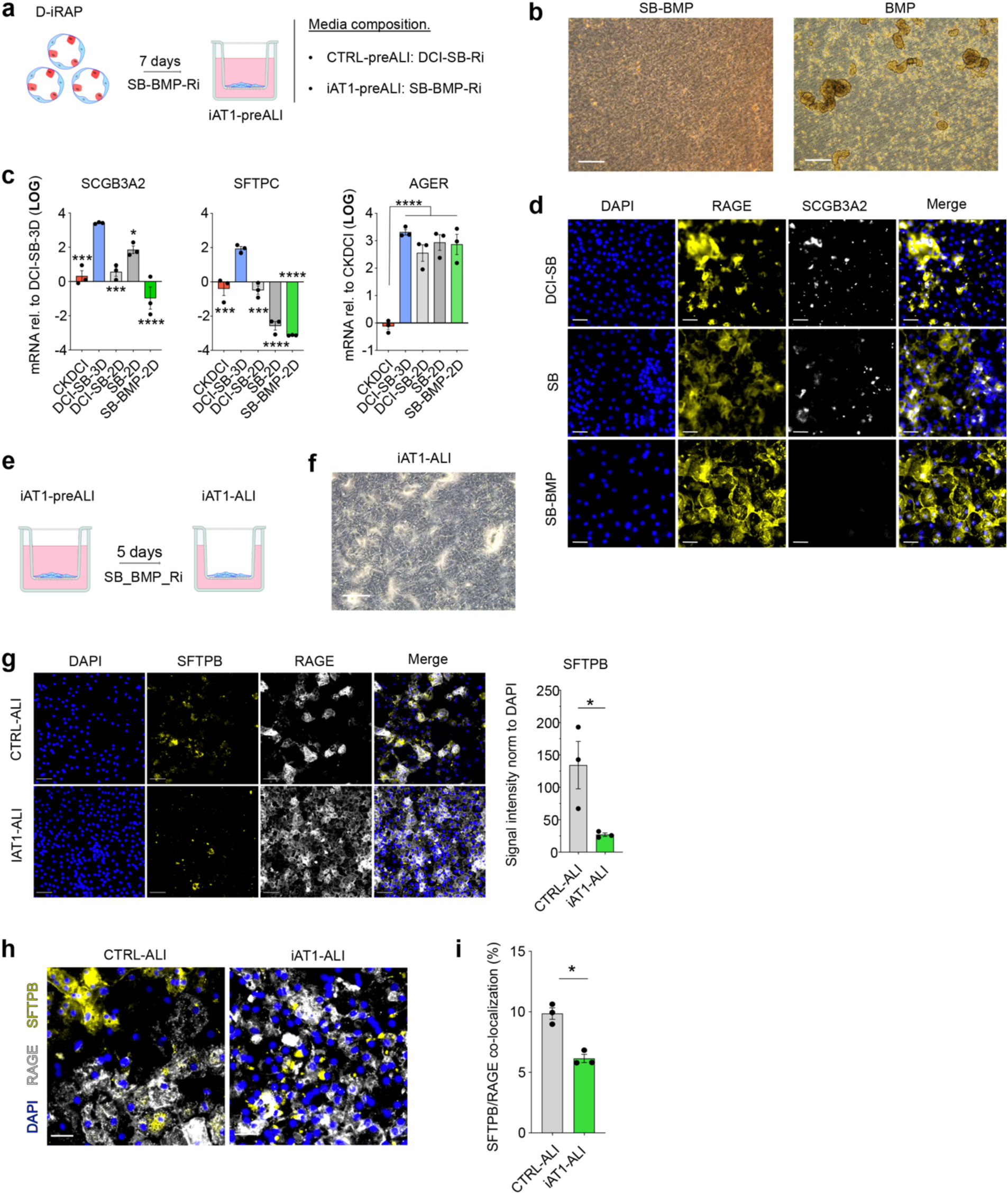
Further characterization of iAT1 differentiation. **a.** Schematic of iAT1 differentiation in 2D submerged culture. **b.** Representative bright field images of 2D submerged cultures in the indicated conditions. Scale bar = 200 µm. **c.** mRNA expression for indicated markers in conditions shown in the X-axis. *p<0.05, **p<0.01, ***p<0.001, ****p<0.0001, one-way ANOVA with Tukey multiple comparison test (n=3). **d.** Representative confocal images after staining for indicated markers in conditions shown on the left. Scale bars = 50 µm. **e**. Schematic of iAT1 differentiation in 2D ALI culture. **f.** Representative bright field image of ALI culture in SB-BMP. Scale bar = 200 µm. **g.** Representative confocal images after staining for indicated markers (left) with quantification (right). Scale bars = 50 µm. *p<0.05, Student’s t-test, n=3. **h.** Representative confocal images showing SFTPB and AGER colocalization (left) with quantification (right). Scale bars = 100 µm. *p<0.05, Student’s t-test, n=3.

**ED8.**
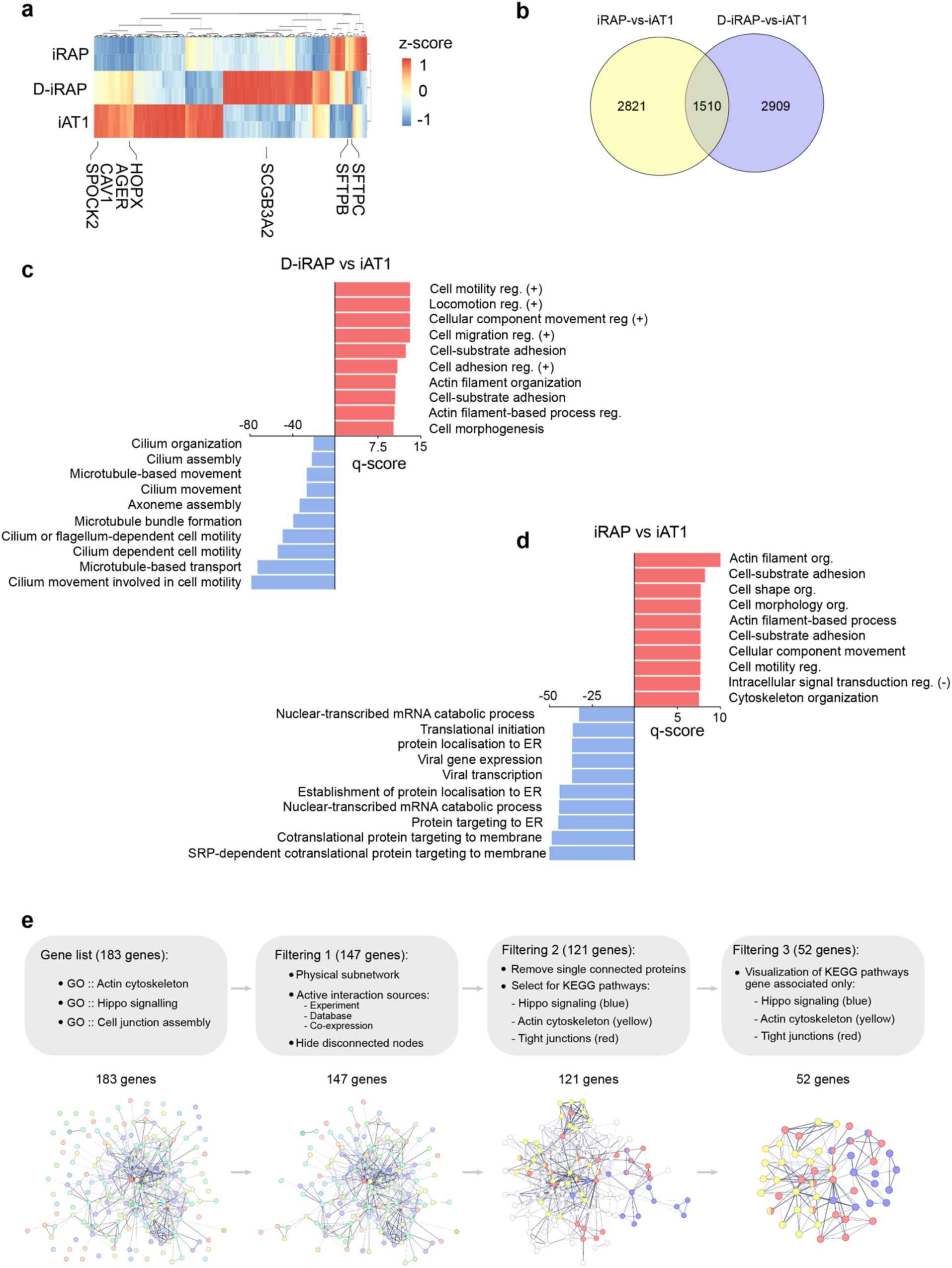
Bulk RNA-seq of iAT1 differentiation. **a.** Heat map representing normalized gene counts throughout iAT1 differentiation. **b.** Venn diagram showing the number of DEGs between the indicated samples. **c.** Top 10 up (red) and down (blue) regulated GO pathways between D-iRAPs and iAT1 cells . **d.** Idem between iRAPs and iAT1 cells. **e.** Protein-protein interaction analysis pipeline analysis.

**ED9.**
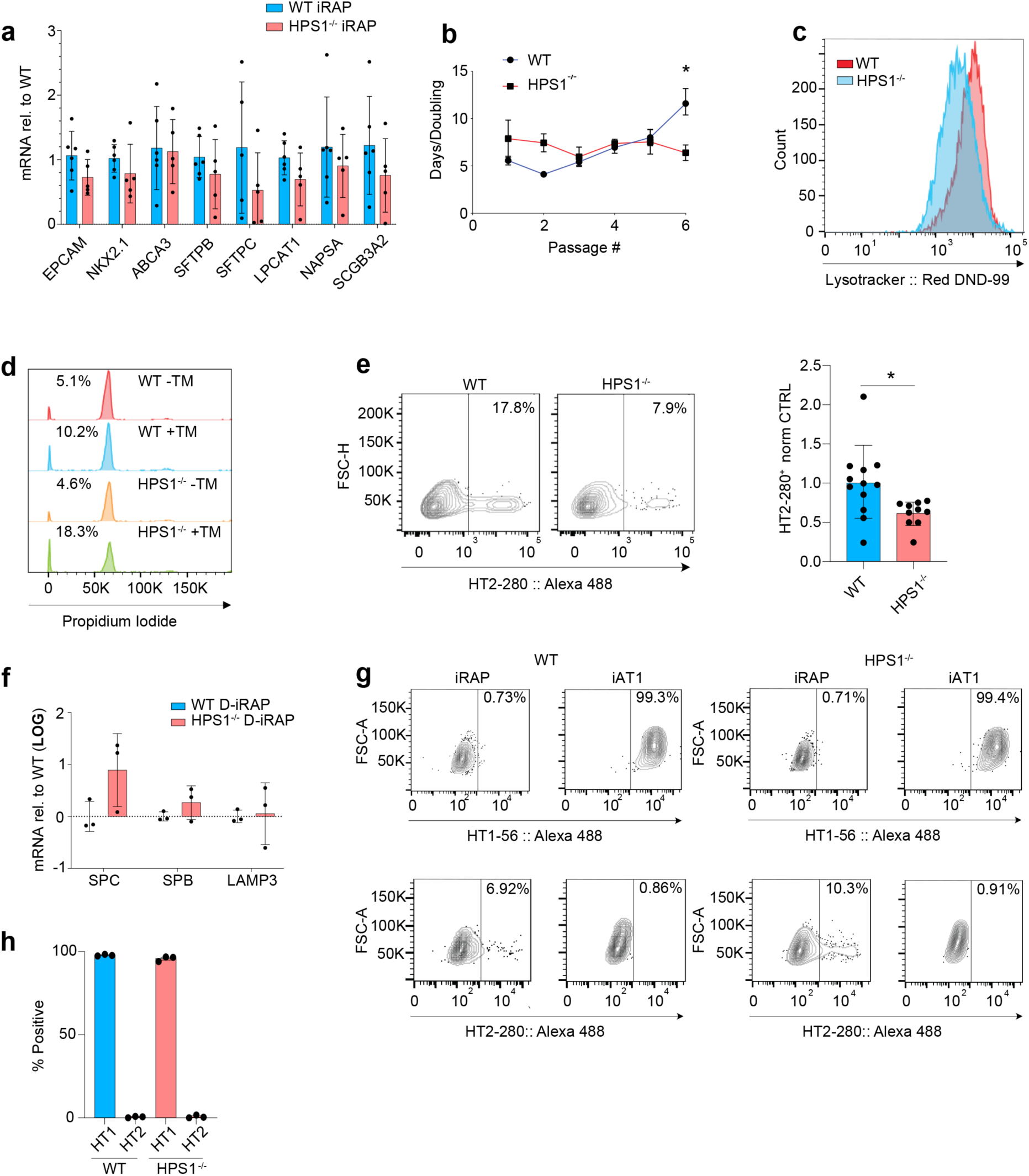
Further characterization of *HSP1^-/-^* iRAPs. **a.** mRNA expression for lung markers in *HPS1*^-/-^ and WT iRAPs. Student’s unpaired t-test, WT n=6, HPS1^-/-^ n=3. **b.** Days per population doubling in WT and HPS1^-/-^ iRAPs. *p<.05, Two-way ANOVA with Sidak’s multiple comparison test, n≥3. **c.** Representative flow cytometry histogram of Lysotracker^®^ uptake in WT and *HSP1^-/-^*iRAPs. **d.** Representative flow cytometry histogram of propidium iodide staining of iRAPs with and without 5 days of 0.1 ng/mL tunicamycin exposure. **e.** Representative flow cytometry contour plot of HT2-280 expression in WT and *HPS1^-/-^* D-iRAPs (left) with quantification of HT2-280^+^ cells in *HSP1^-/-^* D-iRAPs normalized to WT. *p<.05, Student’s unpaired t-test, n≥10. **f.** mRNA expression for indicated markers in *WT* and *HPS1*^-/-^ D-iRAPs compared to iRAPs of the same genotype. ***p<0.001, Two-way ANOVA with Tukey’s multiple comparison, n=3. **g.** Representative flow cytometry plots of HT1-56 and HT2-280 expression in *WT* and *HPS1^-/-^* iAT1 cells. **h.** Quantification of (**g)**.

**Table S1:**
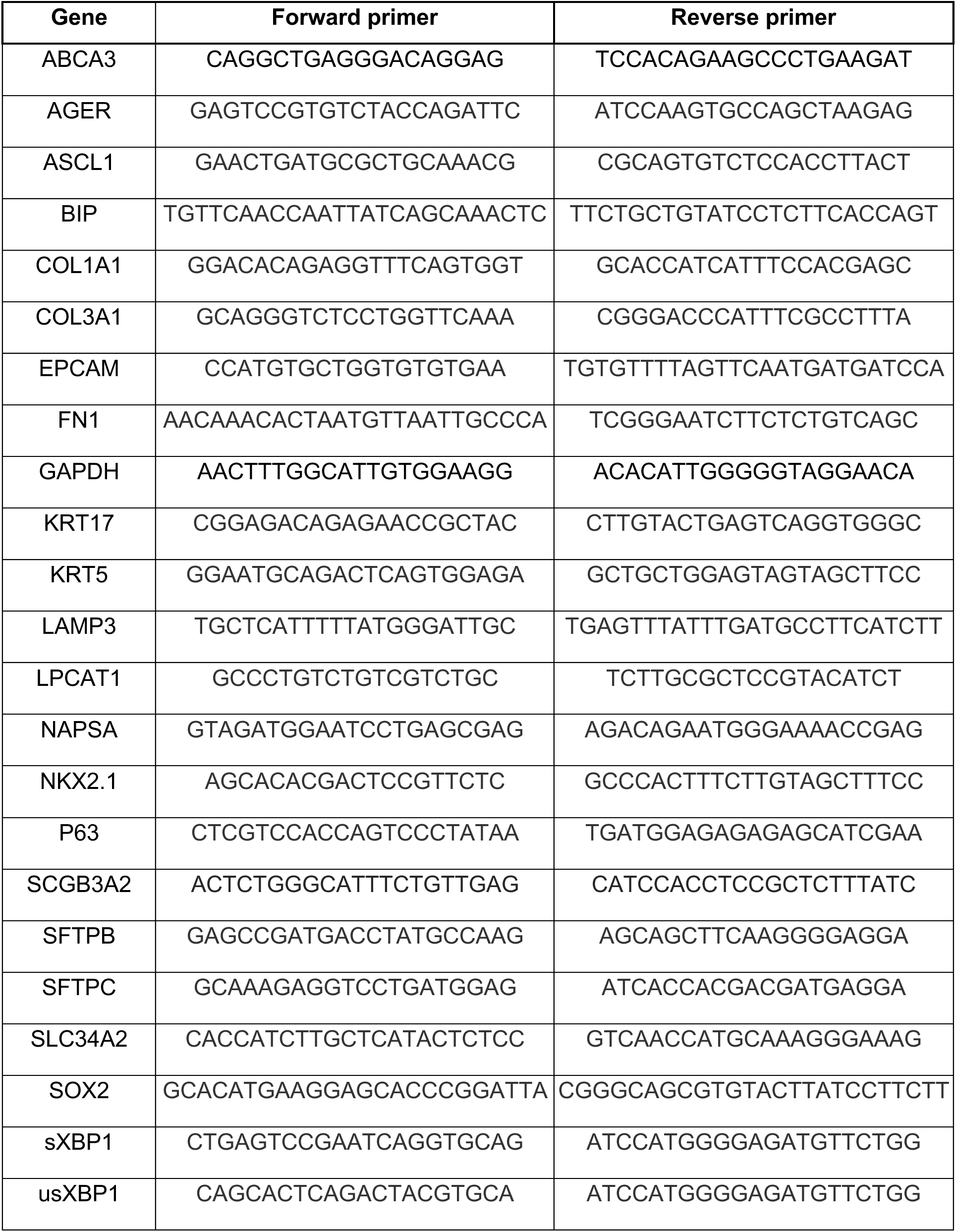
PCR primers.

**Table S2:**
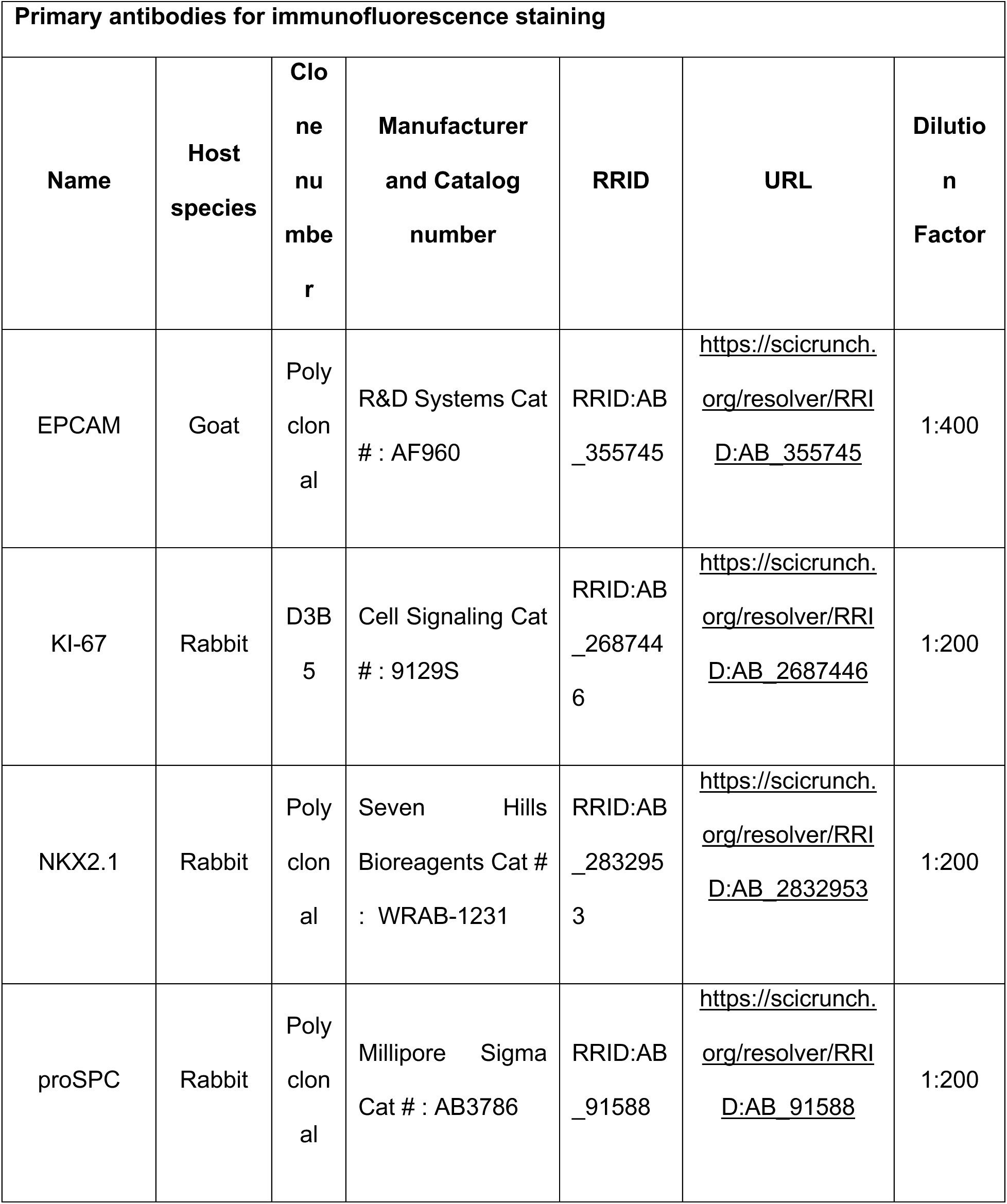
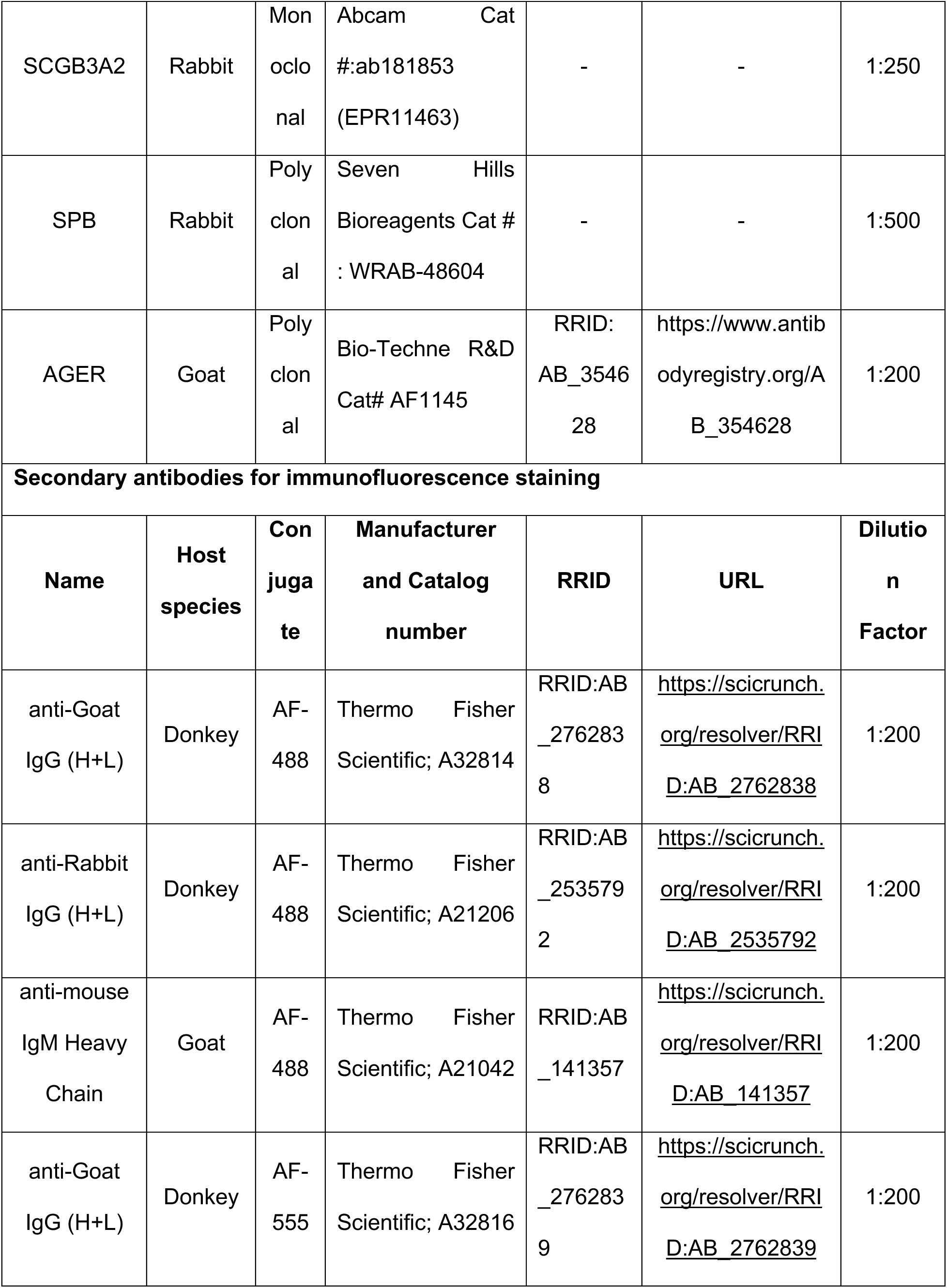
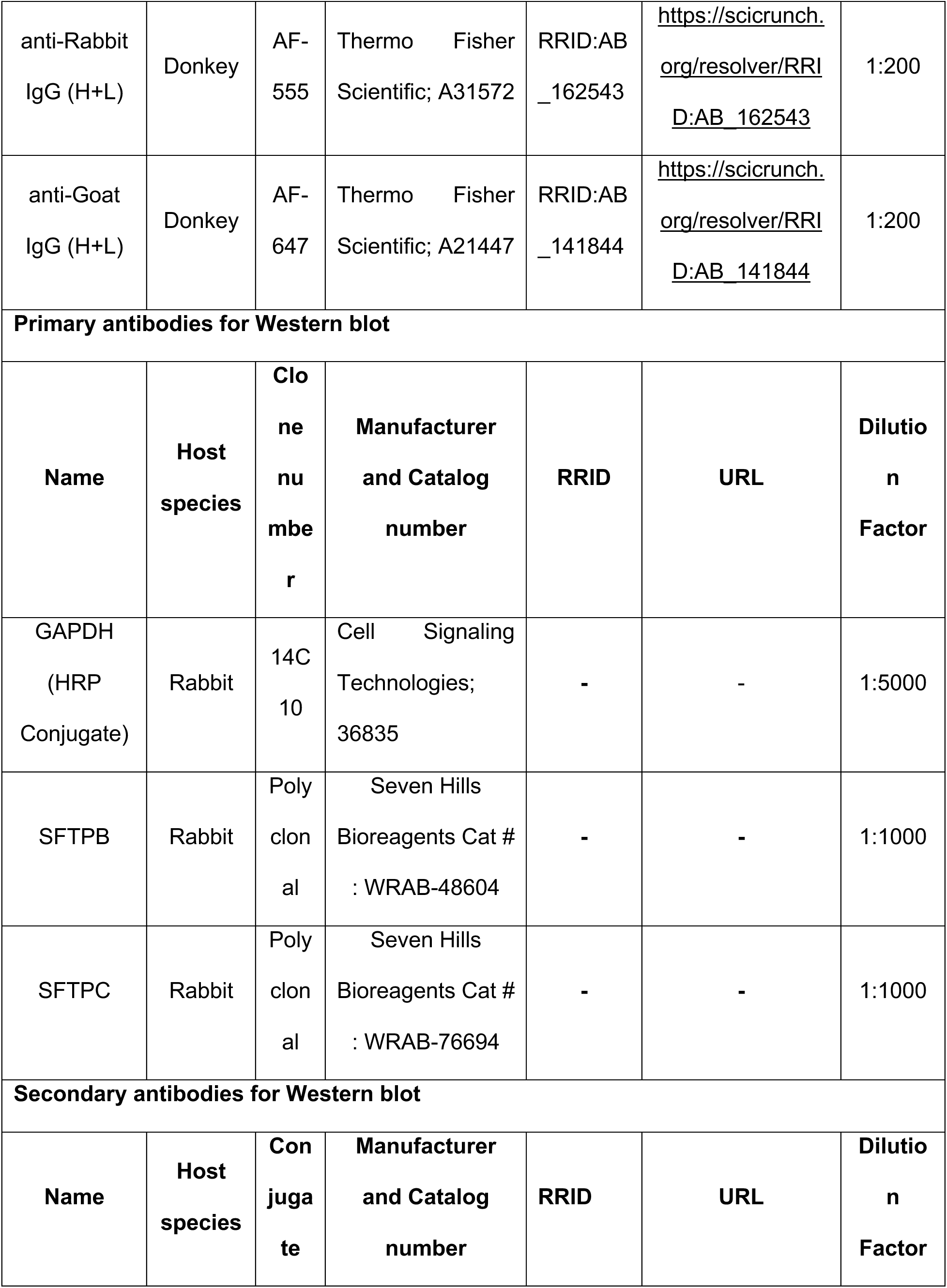
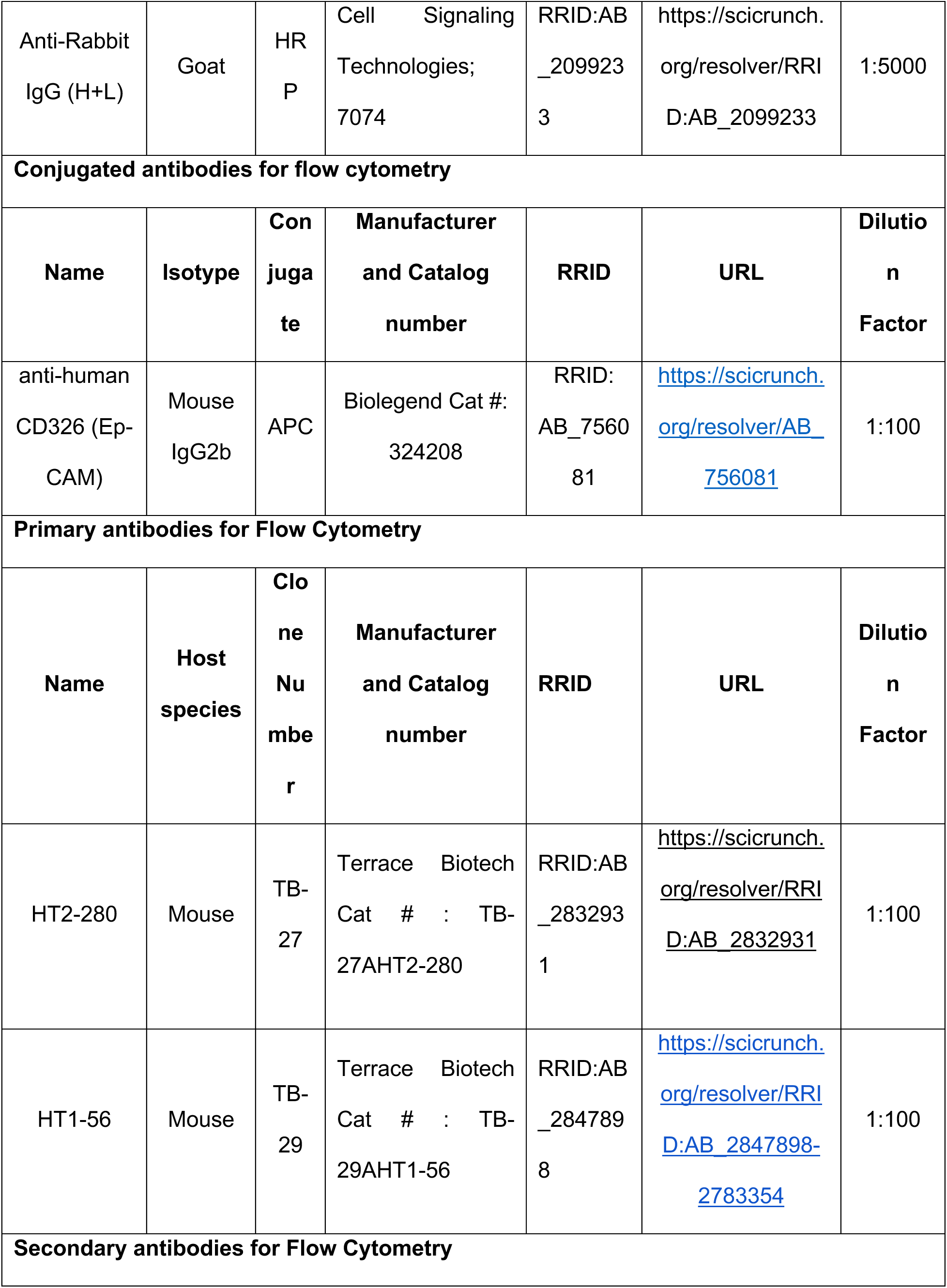
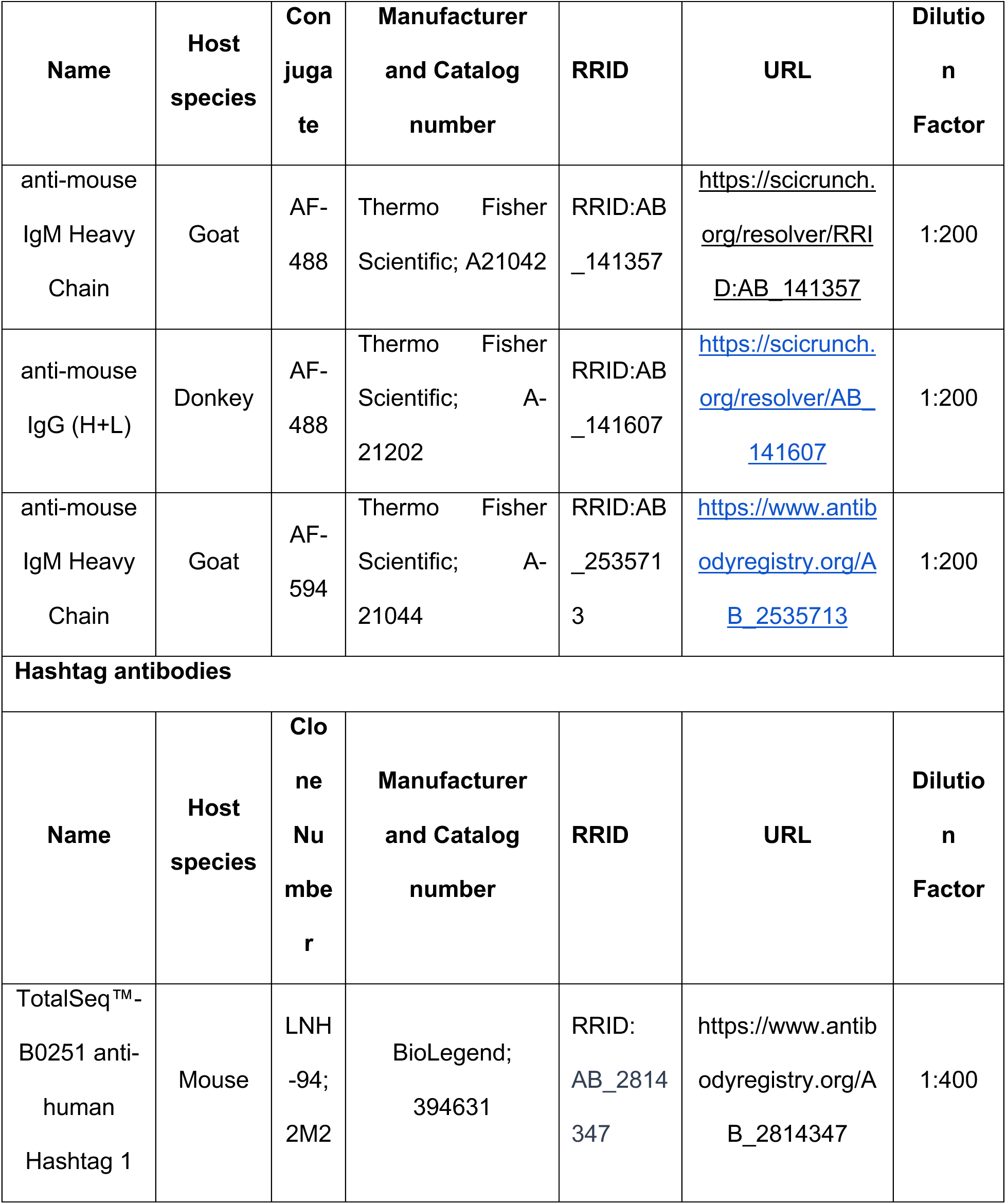
Antibodies.

